# Influence of FTDP-17 mutants on circular Tau RNAs

**DOI:** 10.1101/2023.09.08.556913

**Authors:** Giorgi Margvelani, Justin R. Welden, Andrea Arizaca Maquera, Jennifer E. Van Eyk, Christopher Murray, Sandra C. Miranda Sardon, Stefan Stamm

## Abstract

At least 53 mutations in the microtubule associated protein tau gene (MAPT) have been identified that cause frontotemporal dementia. 47 of these mutations are localized between exons 7 and 13. They could thus affect the formation of circular RNAs (circRNAs) from the MAPT gene that occur through backsplicing from exon 12 to either exon 10 or exon 7. We analyzed representative mutants and found that five FTDP-17 mutations increase the formation of 12➔7 circRNA and three different mutations increase the amount of 12➔10 circRNA. CircRNAs are translated after undergoing adenosine to inosine RNA editing, catalyzed by ADAR enzymes. We found that the interferon induced ADAR1-p150 isoform has the strongest effect on circTau RNA translation. ADAR1-p150 activity had a stronger effect on circTau RNA expression and strongly decreased 12➔7 circRNA, but unexpectedly increased 12➔10 circRNA. In both cases, ADAR-activity strongly promoted translation of circTau RNAs. Unexpectedly, we found that the 12➔7 circTau protein interacts with eukaryotic initiation factor 4B (eIF4B), which is reduced by four FTDP-17 mutations located in the second microtubule domain. These are the first studies of the effect of human mutations on circular RNA formation and translation. They show that point mutations influence circRNA expression levels, likely through changes in the secondary pre-mRNA structures. The effect of the mutations is surpassed by editing of the circular RNAs, leading to their translation. Thus, circular RNAs and their editing status should be considered when analyzing FTDP-17 mutations.

**Highlights:**

- 47/53 known FTDP-17 mutations are located in regions that could influence generation of circular RNAs from the MAPT gene
- Circular Tau RNAs are translated after adenosine to inosine RNA editing, most effectively caused by ADAR1-p150
- FTDP-17 mutations influence both circTau RNA and circTau protein expression levels
- CircTau protein expression levels do not correlate with circTau RNA expression levels
- CircTau proteins bind to eukaryotic initiation factor 4B, which is antagonized by FTDP-17 mutations in exon 10

**Graphic Abstract:** 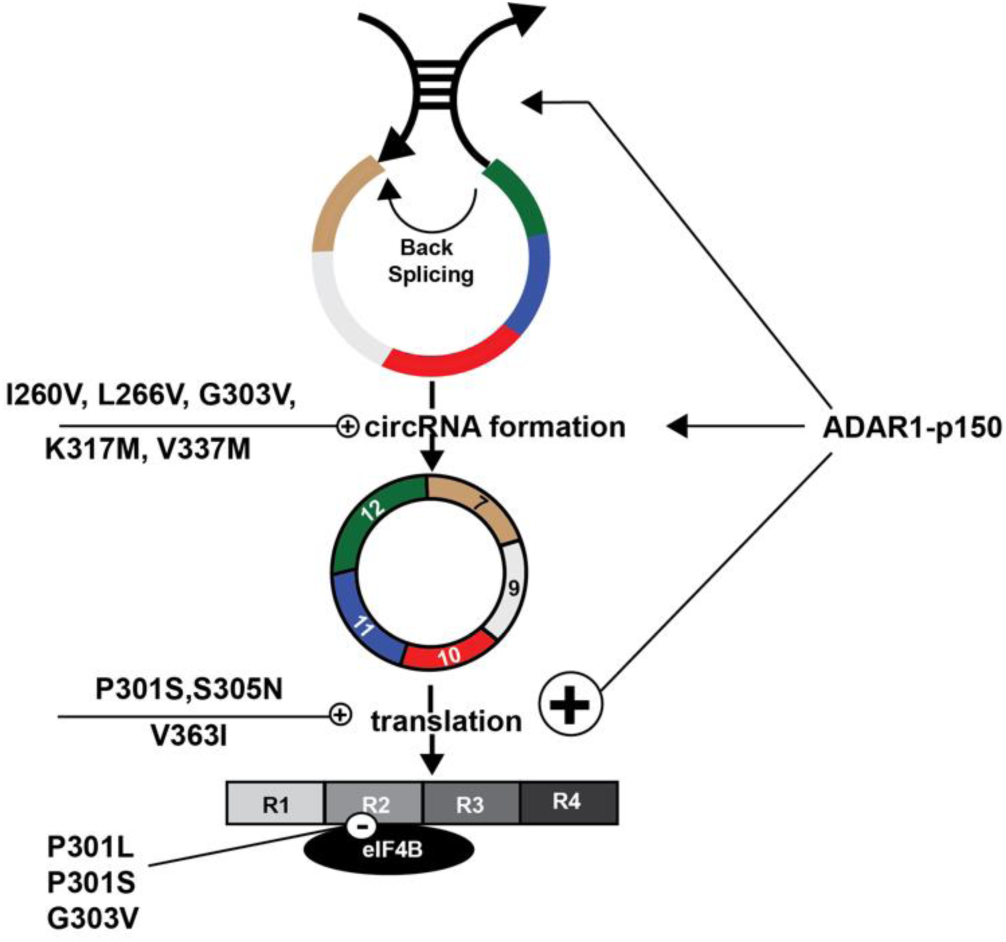

## 1. Introduction

### 1.1 CircTau RNAs

Circular RNAs (circRNAs) are a recently identified class of RNAs, generated through backsplicing of a 5′ splice site to a downstream 3′ splice site [1]. They are expressed in many tissues at low levels, usually less than 1% of their linear counterparts [2,3]. CircRNAs are highly expressed in brain [4–7], are cytosolic and enriched in synaptosomes. Changes in expression of 33 circRNAs have been detected in parietal cortex in Alzheimer’s disease (AD) [8]. When combined with publicly available datasets, 14 circRNAs were associated with Braak stages and explained 31% of the clinical dementia rating, compared to 5% of the APOE4 alleles [8], suggesting a correlation between circRNA expression and AD.

CircRNAs were found to be translated [9–11] from reporter constructs, which was surprising, as circRNAs lack ribosomal entry sites or a 5′ cap to allow translational initiation. We found that adenosine to inosine (A>I) RNA editing strongly promotes translation of circRNAs [12,13], suggesting that inosines could provide a ribosomal entry site. CircRNAs are more stable than linear mRNAs, with a half-life of 2-4 days [14,15], which compares to an average half-life of 4 hrs for mRNAs[16]. Due to intramolecular base pairing, circRNAs form rod-like structures with extended double strand RNA sequences [17], and are thus more prone to undergo modifications by ADAR enzymes (adenine deaminase acting on RNA) that recognize double stranded RNA structures and convert adenosines into inosines [18]. Humans express three ADAR enzymes: ADAR1 expressed in all tissues with an interferon-induced cytosolic (p150) and a constitutive nuclear (p110) isoform; ADAR2 showing weak expression in brain and the catalytic inactive ADAR3 showing strong expression in brain.

### 1.2 FTDP-17 mutations that cause tauopathies predominantly localize within the circTau RNAs

The human microtubule-associated protein tau (MAPT) stabilizes microtubules in the brain [19] through up to four microtubule binding repeats, formed by exons 7-12. Exon 10 encodes the second microtubule-binding repeat and is alternatively spliced in adult human brain that contains a mixture of tau proteins with 3 or 4 microtubule binding repeats [20], which is changed in disease [21–23], especially in several forms of frontotemporal dementia with parkinsonism linked to chromosome 17 (FTDP-17), now called frontotemporal lobar degeneration (FTLD-TAU) [24]. In Alzheimer’s disease and related tauopathies, tau protein mis-folds into intracellular, insoluble protein polymers [25] and intraneuronal tau inclusions (neurofibrillary tangles, or NFTs) that are the pathological feature most strongly associated with cognitive status.

We found that the tau gene generates two circular RNAs [26] that are translated after A>I RNA editing [12]. The circTau proteins promote the aggregation of linear tau protein *in vitro* and in reporter cells [12]. CircTau RNAs are divisible by three and do not contain a stop codon. They thus form multimers of the tau microtubule repeat regions.

At least 53 mutations in the *MAPT* gene cause familial tauopathies have been identified. 38/53 of these mutations are in the circTau RNAs and 9/53 of them are in the introns surrounding exon 10 [24]. Thus 47/53 of all FTLD-TAU mutations could influence properties or formation of circular RNAs. Two FTDP-17 mutations, V337M and K317M strongly increase translation of the start codon-less tau 12➔10 circRNA, suggesting that some FTDP-17 mutations could act through circTau RNAs [12].

Here we analyze the influence of exonic FTDP-17 mutants on circTau RNA and circTau protein formation and expression levels. All circTau RNAs with FTDP-17 mutants were expressed as proteins after undergoing A>I editing. We found that distinct FTDP-17 mutants increase circTau protein and circTau RNA expression levels. There is no correlation between the circRNA amounts and levels of generated proteins, suggesting a predominant translational regulation for the formation of circTau proteins. CircTau protein formation is strongly influenced by ADAR activity that is different between various FTDP-17 mutants. Unexpectedly, we found that the 12➔7 circTau protein binds to eukaryotic initiation factor 4B (eIF4B), which is reduced by FTDP-17 mutants located in the second microtubule binding domain in exon 10. This interaction is specific for the protein products of circular, but not linear Tau RNAs. Our data indicate a new mode of action for some FTDP-17 mutants that could contribute to the pathology of frontotemporal dementia.

## 2. Material and Methods

### 2.1 Transfection

We performed transfection in Human Embryonic Kidney (HEK) 293T cells. 4 µg aliquots of Plasmid DNA were mixed with sterile 150mM 150µl NaCI, in a ratio of 1 µg DNA per 3 µl polyethyleneimine solution (1 µg/µl), (PEI; Polysciences, 24765-1), co transfection were at 1:1 (4 µg:4 µg), for total RNA (1 µg:1 µg). The mixture was incubated at room temperature for 25 min and added to cells, cultured in 150mm dish (Alkali scientific TDN0150) for protein experiment and in 6-well culture plate (VWR, 100062-892) for total RNA experiment and transfected at around 50-60% confluency. Cells were lysed and analyzed after 96h of transfection.

### 2.2 Antibodies, Western Blot and immunoprecipitation

Cells were lysed in low RIPA buffer (150 mM NaCI, 50 mM Tris pH 7,4, 1% NP-40) and boiled for 5 min at 95°C, Lysates were precleared with mouse IgG coupled to 4% agarose beads (Aviva Systems Biology, OOIB00007), 1 mM MgCI_2_ and Benzonase (EDM Millipore 71205-3). Precleared lysates were immunoprecipitated using M2 anti-Flag magnetic beads (Invitrogen) and protein eluted by boiling in loading buffer. Immunoprecipitates and 20 µl lysate aliquots were used for SDS gel electrophoresis and subsequent Western Blot. Proteins were detected using commercially available antibodies: anti-Flag mouse Antibody M2 (1:1000, Sigma-Aldrich M8823) and rabbit anti 4R-tau Antibody (1:1000, Cosmo Bio CAC-TIP-4RT-P01) and anti-calreticulin (EPR3924, Abcam ab92516).

### 2.3 Expression constructs

The DNA constructs were generated using Gibson cloning New England Biolabs [27], and the final constructs were deposited into Addgene, with accession numbers (203365 - 203368, 203371 - 203376, 203378, 203379, 203382 - 203388, 203390, 203391).

### 2.4 RT-PCR and TaqMan probes

The TaqMan assays were performed using a one-step TaqMan probe kit (TaqMan^TM^ RNA-to C_T_^TM^ *1-Step* Kit, Invitrogen). As the TaqMan probe we used 12➔7 Probe FAM–ACCATCAGCCCCCTTTTTATTTCCT-MGBNFQ. For GAPDH loading control, the GAPDH TaqMan Probe, Invitrogen, Hs02786624_g1 was used. Primers used for circtau were 12➔7 set2 F: TCGAAGATTGGGTCCCTGGA and 12➔7 set2 R: TTTTGCTGGAATCCTGGTGG. (**Supplemental Figure 1**).

### 2.5 In vitro transcription-translation and protein pull-down

The TnT quick coupled transcription/translation system (Promega) was used to generate ^35^S-labeled eIF4B. The circ tau 12➔7 WT protein was isolated as described through immunoprecipitation from cellular extracts using 10 µl of M2 anti-Flag magnetic beads per 1 mg of total protein [12]. The beads were resuspended in 1 x TBST buffer and mixed with 20 µl of resuspended beads with 5 µl of each TNT product and brought up to 35 µl with 1 x TBST. The samples were incubated at 4 °C overnight with agitation, followed by 3 x washing with 50 µl 1 x TBST in 100 µM NaCl. Samples were analyzed on PAGE gels and imaged using a Typhoon FLA 9500.

### 2.6 Protein quantification

After protein was transferred to nitrocellulose membrane, the transfer was tested using Ponceau S, followed by Western blot with the appropriate antibodies. The optical density of the signals was measured by ImageJ (Fiji) [28]. The area to be analyzed was marked using an 8-bit grayscale and the intensity for each band was calculated. The relative intensity was determined by dividing the Flag-signal with the calreticulin signal, followed by normalization to GFP signal.

## 3. Results

### 3.1 FTDP-17 mutations influence expression levels of the 12➔7 circTau proteins

To test the effect of FTDP-17 mutants on circTau protein expression, we used circTau expression constructs that we transfected into HEK293T cells. The wild-type construct lacking FTDP-17 mutants has been described previously [12]. It consists of exons 7-13 without the internal introns and is flanked 1384 and 957nt of the natural introns six and thirteen. We introduced selected exonic FTDP17-mutations into this construct and determined protein expression using a 3x Flag tag in exon 7 (**Fig. 1A**). Similar to linear tau protein, circTau proteins are heat stable and as described we used the supernatant of boiled lysates from transfected cells for analysis [12].

**Figure 1:**
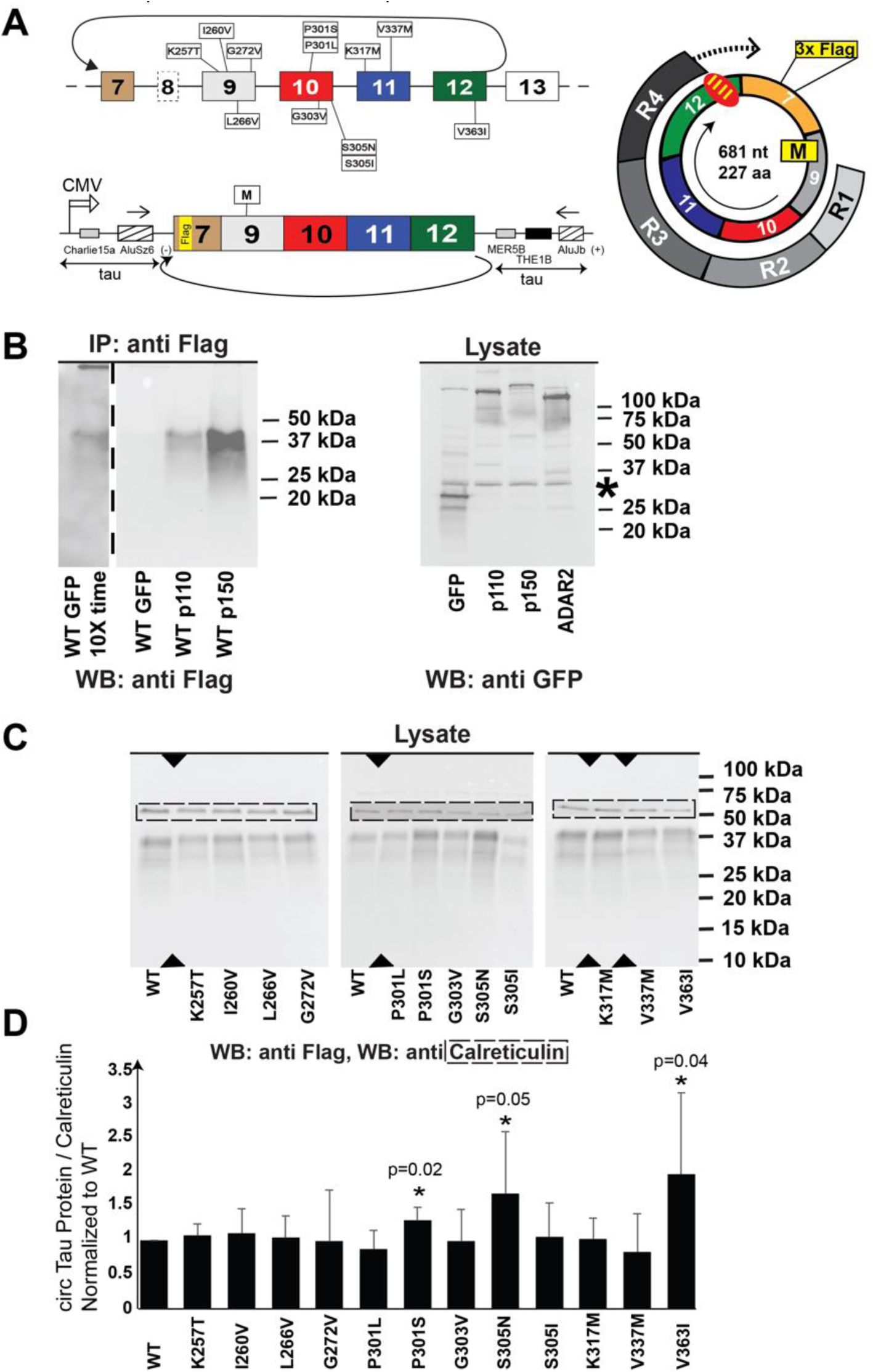
FTDP-17 mutations change expression levels of proteins encoded by the 12૪7 circRNA. **A. Location of selected FTDP-17 mutations in the MAPT gene.** Boxes are exons, lines are introns. The arrows on top indicate the 12➔10 and 12➔7 backsplicing. The FTDP-17 mutations are indicated. The yellow box in exon 9 is the 3x Flag tag used for detection. The structure of the reporter gene is shown schematically underneath and was described in [12], M: single start codon in exon 9. The structure of the circular RNA and its translation product are shown on the right. IIII: stretch of inosines that could serve as a stop signal. R1-R4: microtubule repeat binding domains. **B. Effect of ADAR1-p150 and ADAR1-p110 on circTau-protein expression.** The circTau 12➔7 reporter gene shown in (A) was cotransfected with expression clones for GFP, ADAR1-p150 and ADAR1-p110. CircTau protein was immunoprecipitated using anti-Flag antisera and detected with anti-Flag. The expression of EGFP-tagged ADAR1-p150 and ADAR1-p110 was tested in total cell lysates using Western Blot. Similar volumes of cell lysates were analyzed by western blot using the GFP-tag of the transfected proteins. A star indicates an unspecific band. **C. Effect of FTDP-17 mutants on circTau protein expression.** FTDP-17 mutations were introduced into the 12➔7 circRNA expression construct and were cotransfected with ADAR1-p150. 4 µg expression construct was cotransfected with 4 µg ADAR1-p150 expression construct in 150 mm dishes. Lysates of the transfected cells were separated on PAGE gradient gels and analyzed by Western blot. First, we used an anti-Flag antibody, and the same membrane was reprobed with anti-calreticulin (dotted box). Triangles indicate lanes that were cut out. **D. Quantification of the protein expression.** The ratio of calreticulin signal to Flag-tag signal was calculated and normalized to wild-type. The data represent at least three independent transfection experiments.

We previously reported that circular RNAs are translated after undergoing adenosine to inosine RNA editing, caused by ADAR1 or ADAR2 [12,29]. There are two major isoforms of ADAR1: ADAR1-p150 and ADAR1-p110. They differ in the N-terminal Z-dsRNA binding domain and a nuclear export signal, which are present only in ADAR1-p150 [30]. The two domains are included due to an upstream promoter that is induced by interferons. We thus first compared ADAR1-p150 and ADAR1-p110 for their ability to induce circTau RNA translation by cotransfecting GFP, ADAR1-p110 and ADAR1-p150 expression constructs with circTau wild type expression constructs into HEK293T cells. The presence of ADAR1 activity strongly increased protein abundance and ADAR1-p150 had an about 5-fold stronger effect than ADAR1-p110, even though western blot analysis showed that in the transfection experiments ADAR2 was expressed slightly higher than ADAR1 (**Fig. 1B**). For further experiments, we thus concentrated on the ADAR1-p150 isoform.

The effect of FTDP-17 mutations on circTau protein formation was next tested in cotransfection experiments using expression clones for ADAR1-p150 and the various FTDP-17 mutants. To avoid problems with possible uneven immunoprecipitations due to the FTDP-17 mutations, we exploited the heat stability of circTau proteins and analyzed boiled lysates of the transfected cells. The amount of Flag-tagged circTau proteins was compared with endogenous calreticulin, a protein that is also heat stable [31]. The 12➔7 circTau RNA lacks a stop codon and most likely a stretch of four inosines causes termination of the rolling circle translation. Similar to wild type, the major translation product of circTau proteins harboring the FTDP-17 mutations has a molecular weight of 38 kDa that was seen for all mutants (**Fig. 1C**). When compared to wild type, three mutants, P301S, S305N and V363I showed a statistically significant higher protein expression when compared to wild type (**Fig. 1D**). Our data indicate that three FTDP-17 mutations impact protein expression levels of circTau proteins.

### 3.2 ADAR1-p150 reduces the influence of FTDP-17 mutations on 12➔7 circTau RNA expression

To date, not much is known about the translational mechanism of circular RNAs. To start getting insights into the translational mechanism of circTau RNAs, we first ask whether there is a correlation between the amounts of circTau RNA and the corresponding protein. We devised a TaqMan assay using a custom-made TaqMan probe flanked by PCR-primers in the circRNA (**Fig. 2A**). The TaqMan probe was used with 1 µg of total RNA in a one-step reverse-transcriptase reaction. We used total RNA from cells cotransfected with constructs expressing 12➔7 circTau and GFP. Since TaqMan assays for circular RNAs are novel, we further validated the PCR products after 40 cycles on agarose gels and observed the predicted 216 nt band for all mutants tested (**Supplemental Figure 2A, B**). CircTau RNA was determined from total RNA by calculating the delta ct values of circTau RNA and linear GAPDH. Notably, our assay is sensitive enough to omit the RNase R digestion step used in most circRNA isolation procedures and generated only one band. When using RNase R treated total RNA that is enriched for circular RNAs, we observed multiple bands due to rolling circle reverse transcription (**Supplemental Figure 2C),** which could complicate a TaqMan assay. The 12➔7 circTau RNA signal was normalized to GAPDH mRNA. In contrast to protein expression, several FTDP17-mutants, mainly G303V, I260V and V337M strongly increased circTau RNA expression (**Fig. 2B**).

**Figure 2:**
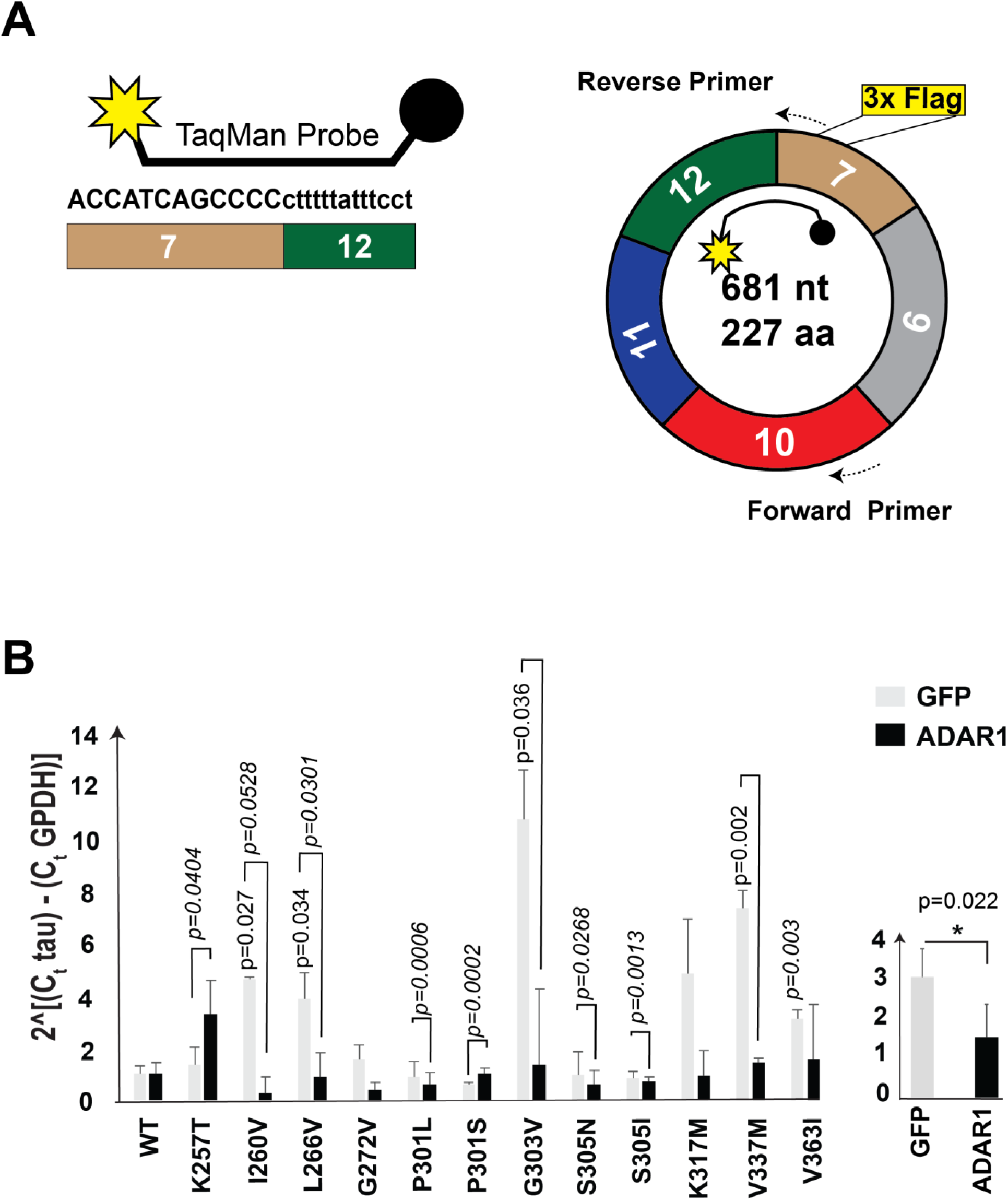
Effect of FTDP-17 mutations and ADAR1-p150 on expression of the 12➔7 circTau RNA. **A.** Sequence and location of the TaqMan probe spanning exons 12 and 7. Star: FAM, circle: MGBNFQ quencher. The location of the primers in the circRNA is schematically shown on right. **B.** Quantification of at least three independent experiments. The 12➔7 circTau signal was normalized to GAPDH. All data were normalized to wild type, which was set to 1. The graph on the right shows the combined delta delta Ct values in the absence or presence of ADAR1-p150.

Unlike the protein, circTau RNA is generated in the absence of ADAR1, allowing us to compare the circTau RNA expression levels in the absence and presence of ADAR1 activity. We thus repeated the experiment using total RNA from cells cotransfected with expression constructs for 12➔7 circTau and ADAR1-p150.

With the exception of K257T, ADAR1-p150 presence reduced circTau RNA expression levels. When all mutants are combined, ADAR1-p150 causes an about three-fold reduction in RNA expression (**Fig. 2B, right**). Thus, some FTDP-17 mutants influence the expression of non-edited circTau RNA. This effect is antagonized by RNA-editing caused by ADAR1-p150 that generally reduces circTau 12➔7 RNA expression.

### 3.3 FTDP-17 mutations influence expression levels of the 12➔10 circTau proteins

The MAPT gene generates two major circRNAs through backsplicing from exon 12 to either exon 7 or exon 10. We thus tested selected mutants in the 12➔10 circTau RNA background, using the expression construct shown in **Figure 3A**. In contrast to the 12➔7 circTau RNA, the 12➔10 circTau RNA lacks a start codon, which can be introduced by RNA editing where an AUA codon is changed to an AUI codon that can initiate translation [12,32]. In addition, the V337M and K317M FTDP-17 mutants introduce start codons and promote translation in the absence of cotransfected ADAR-activity [12] (**Fig 3B**).

**Figure 3:**
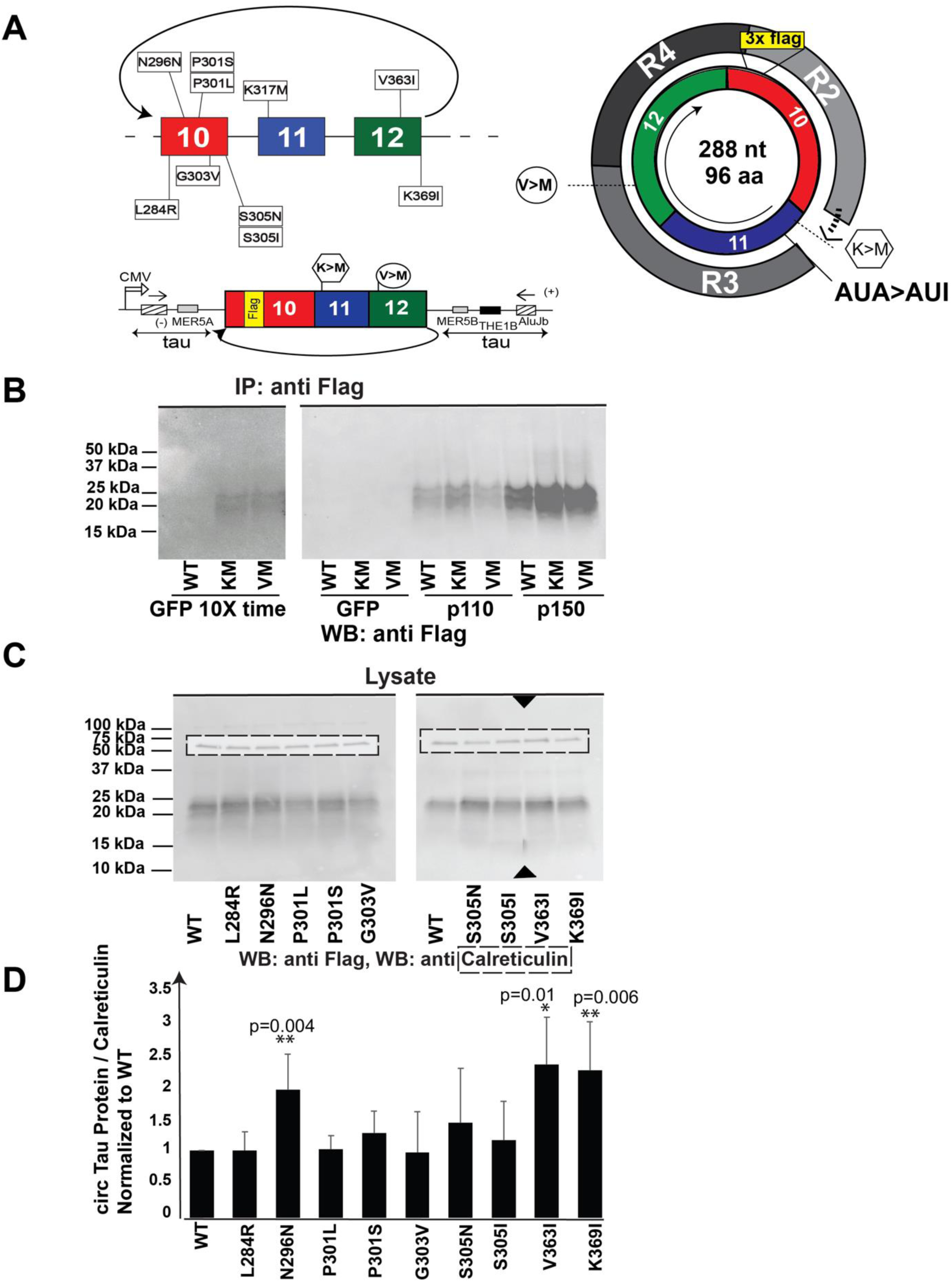
Influence of FTDP-17 mutations on expression levels of proteins encoded by the 12➔10 circRNA. **A.** Schematic overview of the mutations in the 12➔10 circTau RNA. The structure of the expression construct is schematically shown below. The circular RNA is shown on the right. The start codons introduced by the K317M and V337M mutations are indicated. The AUA>AUI editing event generates a start codon that was confirmed by mutagenesis [12]. R2-R3 are tau microtubule binding repeats. **B.** Wild type, K317M and V337M expression constructs were cotransfected with GFP, ADAR1-p110 and ADAR1-p150 expression constructs. circTau protein was isolated using Flag-immunoprecipitation and detected by Flag. **C.** Expression constructs for the indicated mutants were cotransfected with ADAR1-p150 expression constructs in HEK293T cells. Lysates were analyzed by Western blot using Flag and calreticulin antibodies. The calreticulin exposure time is 1/10 of the Flag-tag exposure time. **D.** Quantification of the Flag to calreticulin signal. The ratio was set to 1 for the wild type. At least three independent transfections were analyzed.

We first compared ADAR1-p150 and ADAR1-p110 for their ability to promote circTau RNA translation, using the wild-type, the K317M and V337M mutations. As we previously reported, in the absence of ADAR-activity there is hardly any protein expression detectable from the wild-type, likely due to the absence of a start codon. However, ADAR-activity strongly promoted circTau protein expression. Similar to the 12➔7 circRNAs, ADAR1-p150 has a much stronger effect than ADAR1-p110 (**Figure 3B**). In the absence of cotransfected ADAR activity, the K317M and V337M mutations result in weak protein formation, which is strongly increased by ADAR1. As shown in Figure 3B, ADAR-p150 promotes the translation of the start codon-less 12➔10 circRNA stronger than the introduction of a start codon.

We next tested selected mutants for their influence on 12➔10 circTau protein formation by cotransfecting their expression clones with ADAR1-p150 expression clones in HEK293T cells. All mutants resulted in a protein with the same size as the wild type. Similar to the 12➔7 tau circRNA, we quantified the expressed 12➔10 circTau proteins by comparing them to calreticulin in boiled lysates. As in the 12➔7 background, the mutant V363I had a significant higher expression than wild-type (**Fig. 3D**). Conversely, P301S and S305N increased expression from the 12➔7 background but had no effect on the 12➔10 background. We conclude that one mutant, V363I had a similar enhancing effect on circTau protein expression, but in general the effect of FTDP-17 mutations on protein expression depends on their location in a 12➔7 or 12➔10 circular RNA background.

### 3.4 ADAR1-p150 increases expression levels of the 12->10 circTau RNA

To gain further insight into the translational mechanism of circTau RNAs, we measured the RNA levels of selected FTDP-17 mutations in the 12➔10 background. Several designs of TaqMan probes did not work and we thus performed qPCR using SYBR green (**Fig. 4A**) with the primers indicated in **Supplementary** Figure 3. Like the qPCR analysis of the 12➔7 circTau RNA, we used total RNA from transfected cells and omitted the RNase R treatment step. As shown in **Supplemental Figure 4**, the qPCR resulted in only one band, making the detection with SYBR green possible. The signal from circTau RNA was normalized to linear GAPDH mRNA, measured in a TaqMan assay.

**Figure 4:**
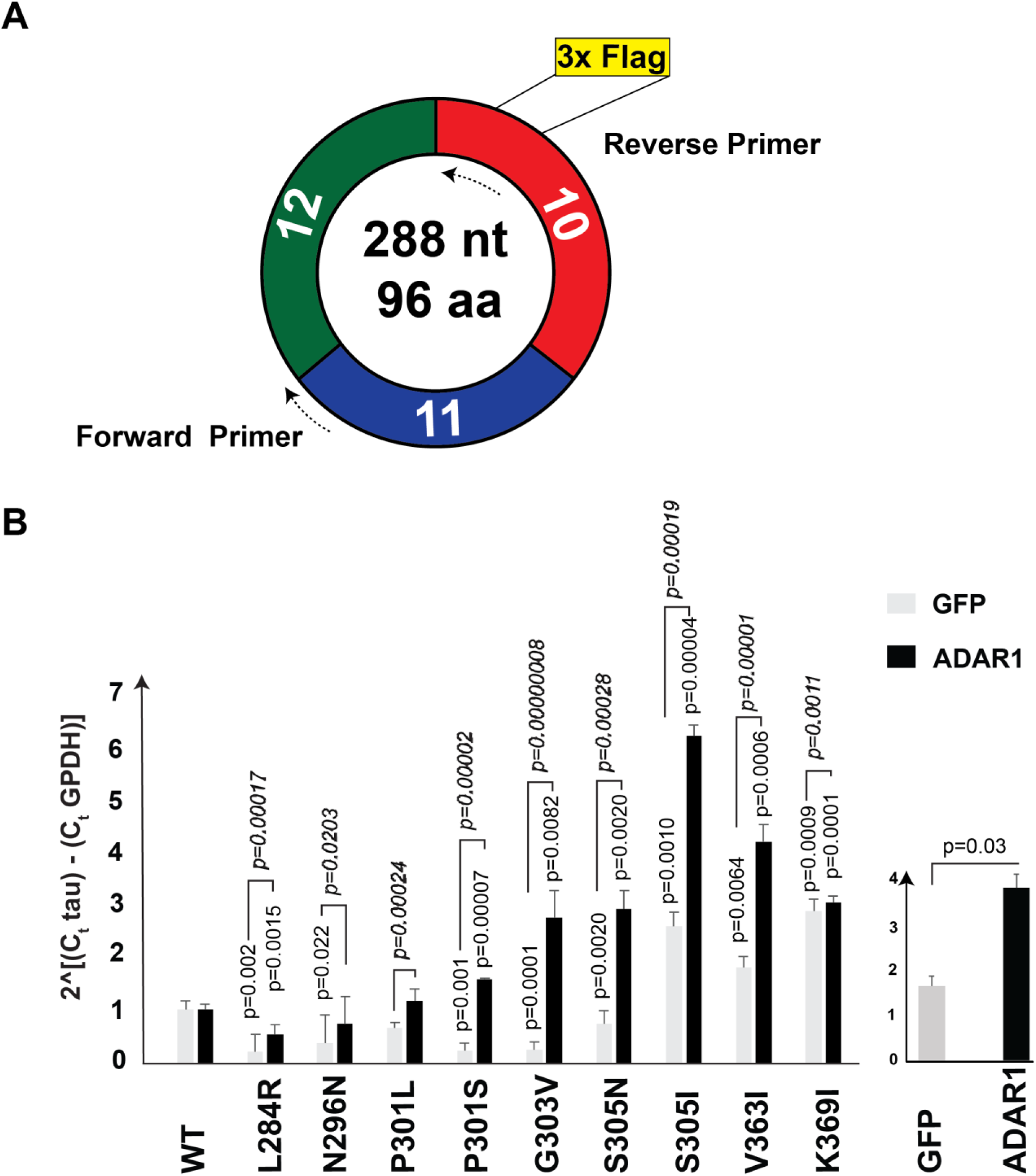
Effect of FTDP-17 mutations and ADAR1-p150 on expression of the 12➔ 10 circTau RNA. **A.** Location of the primers for SYBR green assay in the 12➔10 circTau RNA. **B.** Quantification of the 12➔10 circTau RNA expression of FTDP-17 mutants, which were compared to GAPDH and compared to wild-type, set to 1.

In the absence of ADAR1 activity, the FTDP-17 mutations S305I, V363I and K369I strongly increased 12➔10 circTau RNA expression compared to wild-type. Upon cotransfection of ADAR1-p150, we observed strong expression of five mutants: P301S, G303V, S305N, S305I, and V363I (**Fig. 4B**). The change in 12➔10 circRNA levels is not reflected on the protein level that showed uniform expression (**Fig. 3**), indicating that 12➔10 circTau protein formation is mainly regulated on the translational level. Thus, ADAR1-p150 has different effects on circRNA expression, depending on the circRNA sequence and possibly flanking introns.

### 3.5 Effect of ADAR1-p150, ADAR2 and ADAR3 on translation of FTDP-17 mutants

FTDP-17 mutations vary in clinical presentation, age of onset [33] and their effect on the assembly of tau into filaments [34]. Likewise, ADAR-enzymes are not uniformly expressed throughout the brain. ADAR1 and ADAR2 are expressed in neurons, but not in astrocytes throughout the brain. ADAR1 expression is lowest in cortex, choroid plexus and hippocampus, whereas ADAR2 shows lowest expression in cortex and hippocampus and high expression in thalamus [35]. The catalytic inactive variant ADAR3 is expressed only in brain. We thus asked whether FTDP-17 mutants respond differently to ADAR-1, -2 and -3 and tested eight mutants.

The same amounts of expression clones for these mutants in the 12➔7 circTau background were cotransfected with ADAR expression clones and protein expression in boiled lysates was normalized to calreticulin and quantified. For all mutants tested, ADAR1-p150 had a stronger effect than ADAR2 (**Figure 5A, B**). In addition, three mutants P301L, P301S, and K317M increased the translatability by ADAR2. For most mutants, we saw a statistically significant increase of protein with the catalytic inactive variant ADAR3 when compared to the GFP control. Similar results were observed, when we analyzed the transfection data with an anti-tau antiserum (**Supplemental Figure 5**). The slight increase of circTau protein due to ADAR3 was surprising, but we previously observed it using wild type constructs and also found slight changes in circTau RNA editing [12]. We speculate that ADAR3 transfection increases ADAR1 or ADAR2 activity due to an autoregulatory effect and/or heterodimerization between the ADAR enzymes. The data show that ADAR1-p150 had the strongest effect on translation of circTau RNAs harboring FTDP-17 mutants, and three mutants increase the translation caused by ADAR2.

**Figure 5:**
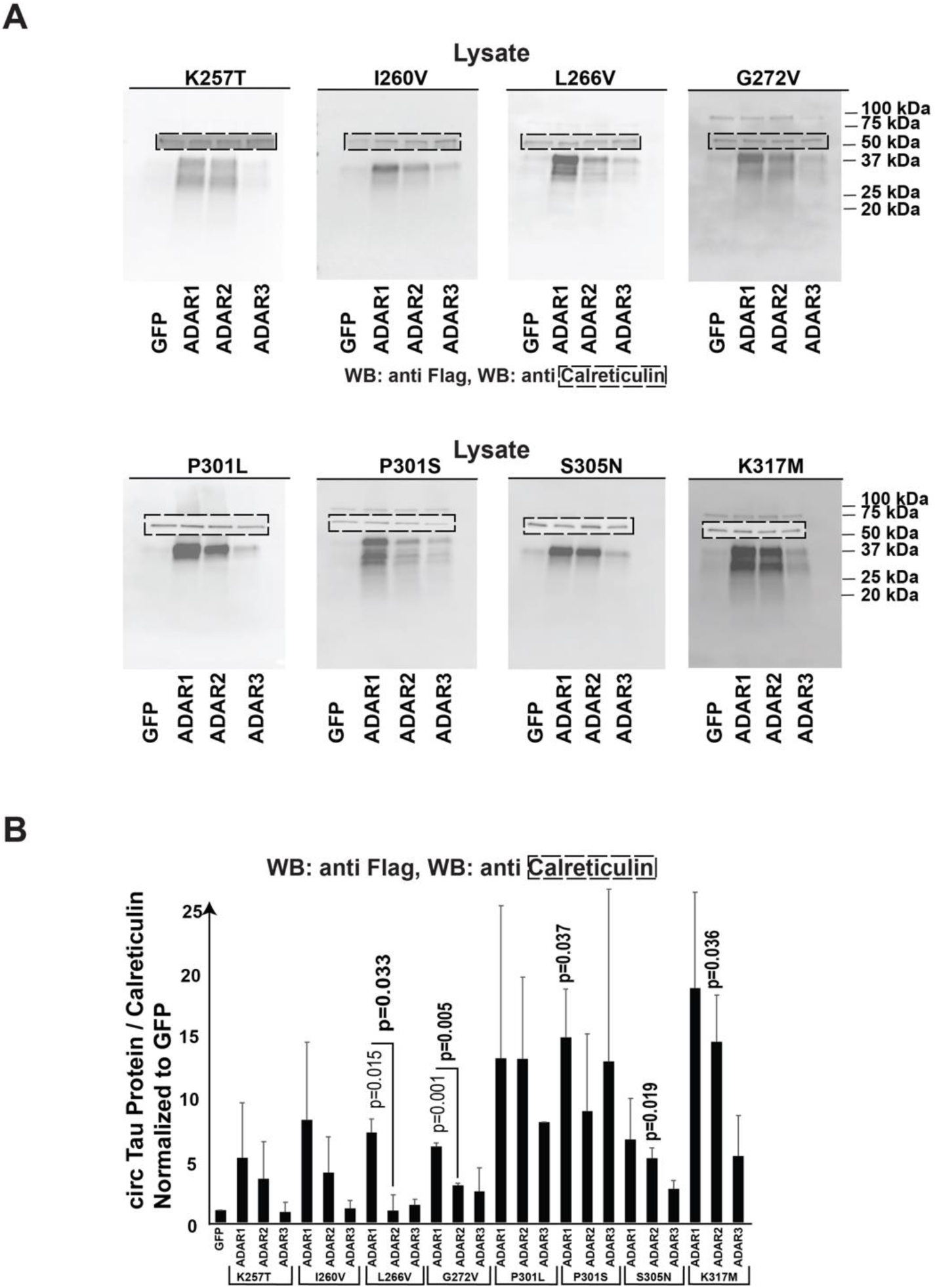
Response of selected FTDP-17 mutants on ADAR1-p150, ADAR2 and ADAR3. **A.** Effect of ADAR1-p150, ADAR2 and ADAR3 on FTDP-17 mutants. Expression clones of the FTDP-17 mutants in the 12➔7 circTau RNA background were cotransfected with expression clones for equal amounts of EGFP, ADAR1-p150, ADAR2 and ADAR3. Equal volumes of boiled lysates were analyzed by Western blot and the blots were re-tested with anti-calreticulin (dotted boxes). **B.** Quantification of three independent experiments. The ratio of calreticulin to circTau-protein of three independent experiments was calculated and normalized to wild-type. bold p-values indicate the significance between GFP and ADAR1-dependent expression and regular font p-values indicate the significance between wild-type and mutants.

### 3.6 Proteins made from the 12➔7 circTau RNA interact with eukaryotic initiation factor 4B

When validating the circTau proteins through mass-spectrometry, we detected interacting proteins in the immunoprecipitates that were generated from boiled lysates (**Supplemental Figure 6**). Other than tau, eukaryotic initiation factor 4B (eIF4B) was the most abundant interacting protein. This finding was unexpected, as binding from eIF4B to the linear tau protein has not been reported.

We first confirmed the specific interaction between 12➔7 circTau protein and endogenous eIF4B in immunoprecipitation experiments. To directly test a possible binding to linear tau protein, we expressed a linear 3xFlag-0N4R construct [12], as well as the 12➔7 and 12➔10 tau circRNA expression constructs in HEK293T cells. All proteins were immunoprecipitated from heat-treated lysates using anti Flag magnetic beads. We first confirmed the expression of the expected proteins by testing the immunoprecipitates with anti Flag antibodies. Using the same volumes of cell lysates, we found that linear 3xFlag-0N4R protein is expressed about 10-fold stronger than the circular proteins (**Fig. 6A, left**). Next, we tested the immunoprecipitates with an anti-eIF4B antibody, which showed presence of eIF4B in the immunoprecipitates of the circular 12➔7 circRNA encoded protein (**Figure 6A, right**), validating the interaction we observed in mass-spectrometry. The interaction with the 12➔10 circRNA encoded protein was significantly weaker. Importantly, we could not detect an interaction between linear tau protein and eIF4B (**Fig. 6A, right**), even though the linear tau protein was expressed at much higher levels. The data suggest that eIF4B interacts predominately with the protein products of the 12➔7 circTau RNAs. The 12➔7 circTau protein is highly similar to a fragment of the linear 3xFlag-0N4R protein and differs only in the exon junction region. Thus, the lack of interaction between eIF4B and 3xFlag-0N4R protein thus suggests a highly specific interaction between the 12➔7 circTau encoded protein and eIF4B.

**Figure 6:**
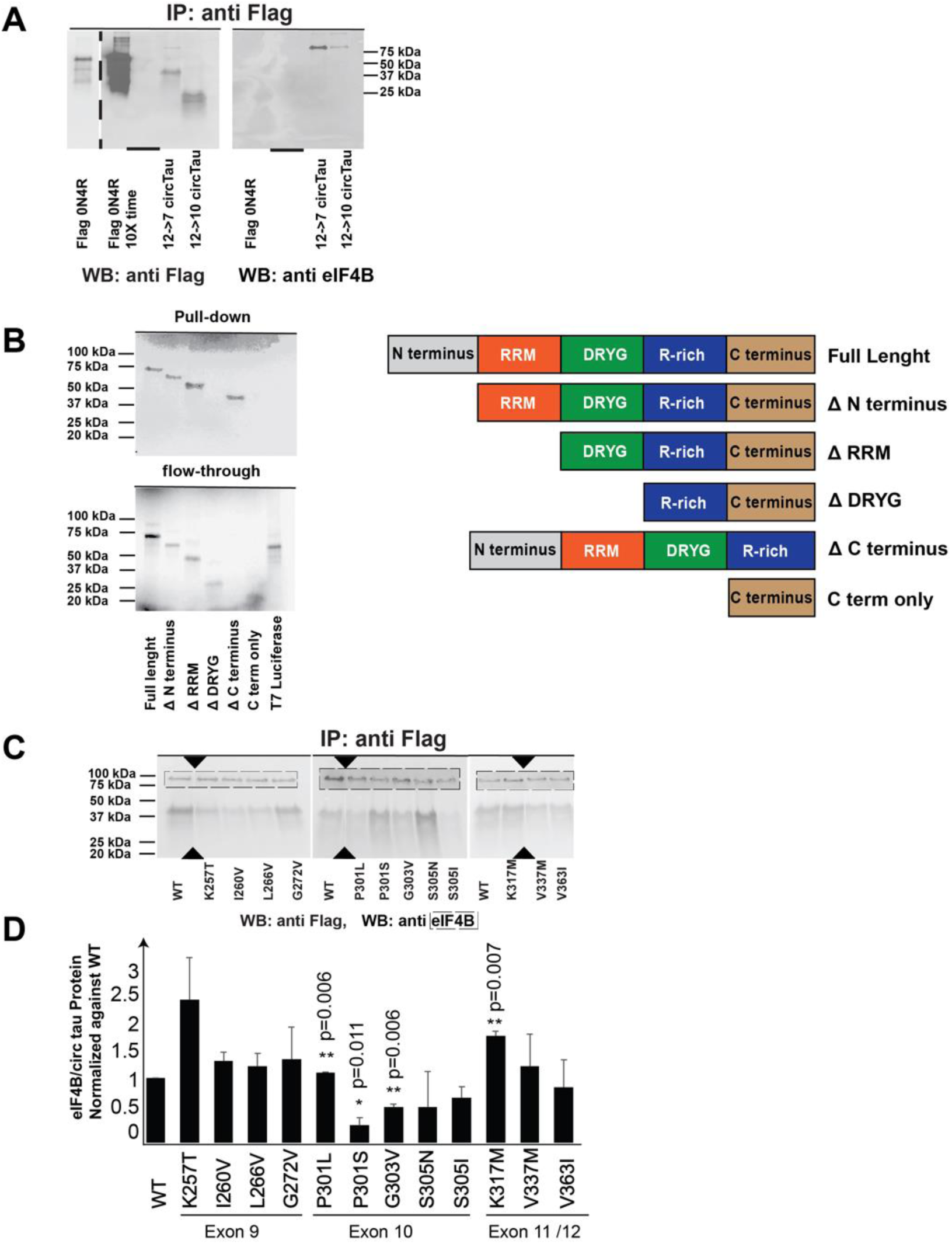
circTau 12➔7 protein binds to eIF4B. **A.** Immunoprecipitation of endogenous eIF4B and linear 0N4R tau and circTau proteins. Clones expressing linear tau 0N4R, as well as 12➔7 circTau and 12➔10 circTau clones were transfected into HEK293T cells and immunoprecipitated with anti-Flag. The immunoprecipitates were tested with anti-Flag and anti-eIF4B. The lane for Flag-0N4R tau is shown with short exposure on the left. “-” empty lane to account for the strong linear tau signal. **B.** Mapping of eIF4B interaction domain. The domain structure for eIF4B is shown on the right. circTau 12➔7 protein was bound to agarose and incubated with eIF4B deletion variants made in reticulate lysates. Bound proteins (top) and flow-through (bottom) were analyzed by PAGE, using the radioactive ^35^S label for detection. **C.** Interaction of eIF4B and 12➔7 circTau proteins. The FTDP-17 12➔7 circTau mutants indicated were expressed in HEK293T cells and immunoprecipitated with anti-Flag. The immunoprecipitates were first analyzed with anti-Flag for circTau proteins and with eIF4B for endogenous eIF4B binding. Triangles indicate removal of lanes from the gel. **D.** Quantification of the eIF4B – circTau protein interaction from three independent experiments. The ratio of eIF4B/circTau protein signal was determined and normalized to wild type.

EIF4B consists of several domains involved in interactions with various proteins (**Fig. 6B**). The N-terminal domain binds to poly A binding protein [36], the RNA recognition motif and an arginine-rich domain that binds to RNA [37], the C-terminal that binds to eIF4A [38], and the DRYG repeat domain binds to eIF3A and aids in eIF4B dimerization [39].

Given the propensity of eIF4B’s domains to interact with different proteins, we hypothesized that similarly eIF4B interacts with 12➔7 circTau protein using a specific domain. To determine this interaction site, we expressed ^35^S-labled eIF4B variants in reticulocyte lysates and incubated 12➔7 circTau wild-type protein bound to agarose through the Flag tag. We analyzed flowthroughs and bound protein, which were visualized using the radioactive label. We observed eIF4B lacking the DRYG domain was absent in the interaction matrix, but present in the flowthrough. All other constructs containing a DRYG domain showed binding to the 12➔7 circTau protein-resin (**Fig. 6B**). These findings indicate that the DRYG domain mediates the interaction between eIF4B and circTau proteins.

### 3.7 FTDP-17 mutations located in exon 10 reduce the interaction between eIF4B and 12➔7 circTau protein

We next tested the interaction between FTDP-17 mutants in the 12➔7 circTau background and eIF4B. The mutant 12➔7 circTau proteins were overexpressed in HEK293T cells and immunoprecipitated from heat treated lysates using anti-Flag. The Flag-tagged 12➔7 circTau protein and endogenous eIF4B were detected in the immunoprecipitates. When normalizing the signal of eIF4B to the 12➔7 circTau signal in the immunoprecipitates, we saw a reduction of eIF4B binding with the mutants P301L, P301S, and G303V, located in exon 10 (**Fig. 6C, D**). Our immunoprecipitation experiments (**Fig. 6A**) indicate that eIF4B binds much weaker to the 12➔10 circTau protein. We tested the influence of selected mutations in the 12➔10 circTau background. Compared to wild-type, we observed a reduction for eIF4B binding for L284R, G303V and P301L, three FTDP-17 mutations located in exon 10 (**Supplemental Figure 7**). All mutations affecting eIF4B binding are located in the second microtubule binding domain in exon 10 suggesting that this region of the circTau protein participates in binding to eIF4B.

## 4. Discussion

The effect of FTDP-17 mutations on the linear tau protein have been extensively studied. Mutants can be subdivided into two groups: those that change alternative exon 10 usage and secondly mutants that act on the protein level [20,24,34,40,41]. In addition to proteins made from mRNA, the human MAPT-gene generates circular RNAs [26] that are translated [12] after some of their adenosine residues are enzymatically converted into inosines due to A>I RNA editing. As 47 of the 53 FTDP-17 mutations are located in regions that could contribute to circTau RNA generation, we studied for the first time the effect of FTDP-17 mutations on circTau RNA formation and translation.

### 4.1 Circular RNAs can be analyzed by TaqMan probes

Not much is known about the editing-dependent translation of circRNAs and we thus first set up a TaqMan assay to quantify circRNA expression levels. In general, the primer options for circRNAs are limited and in the circTau case design programs do not find optimal TaqMan sequences. We thus tested various TaqMan probes against the unique backsplice junction and initially followed the rules for linear RNAs, i.e., a length of 16-30 nt, the melting points about 10-15 °C higher than the PCR primers and a GC-content of 30-80%. We found that the successful TaqMan probe worked with a melting temperature of 5 °C higher than the amplification primers. In our design we avoided a run of G-nucleotides present in the sense strand by using the reverse complement. We omitted the RNase R digestion step normally used to circRNA isolation [15,42]. RNase R treatment resulted in multiple bands in the TaqMan-RT-PCR, probably as a result of priming with RNA fragments and/or rolling circle reverse transcription that is known to generate a ‘ladder’ of PCR amplicons from circular RNAs [43]. Our 12➔7 circTau TaqMan assay generates only one amplicon and thus allows to measure 12➔7 circTau RNA.

### 4.2 ADAR1-p150 has the strongest effect on circRNA translation

Using RNAseq, we previously measured RNA editing of circTau RNAs in HEK293T cells and found very low editing levels in the absence of cotransfected ADAR activity, that are most likely due to the endogenous ADAR1 and ADAR2 present in HEK293T cells [12]. For all FTDP-17 mutants tested, translation of the circTau RNAs is strongly promoted by ADAR activity, provided in our experimental system by cotransfection of ADAR1 or ADAR2 expression clones. The ADAR1 gene generates two major isoforms: similar to ADAR2, the constitutive ADAR1-p110 isoform is predominantly localized in the nucleus. In contrast, the interferon-induced ADAR1-p150 isoform is predominantly cytosolic [18,44] as in addition to the N-terminal Z-alpha domain it contains a nuclear export sequence. The Z-alpha domain is a dsRNA binding domain that binds to Z-RNAs [45]. It remains to be determined whether circRNAs predominantly adopt the Z-RNA conformation making them preferred substrates for ADAR1-p150. As ADAR1-p150 is induced by interferons, the translation of circRNAs could be a direct consequence of released interleukins and thus relevant for Alzheimer’s diseases that is characterized by inflammation [46–48]. We found that when compared to other mutants, P301L, P301S and K317M, increased the protein expression in the presence of ADAR2, which could indicate that their translation is independent from an interferon-induced induction of ADAR1-p150. It is remarkable that single point mutations in the circular RNA have a strong effect on ADAR2 activity, which could be due to a change in RNA secondary structure and/or generation of an optimized ADAR2 recognition site.

### 4.3 No correlation between circRNA and circProtein expression levels

We expressed circTau RNAs in HEK293T cells by transfecting expression constructs harboring FTDP-17 mutants and measured both circTau protein and circTau RNA expression levels. We observed differences in circTau RNA expression levels for some FTDP-17 mutants. Since all circTau expression constructs were driven by the same promoter in an identical background, the changes are unlikely to be due to a transcriptional effect. A more probable explanation is that mutants interfere with circRNA formation by changing the pre-mRNA structure. ADAR1-p150 coexpression strongly decreases 12➔7 circTau expression, and strongly promotes 12➔7 circTau translation. Some FTDP-17 mutants cause differences in edited 12➔7 circTau RNA expression that are not reflected on the protein level. CircRNA translation is likely mainly regulated through translational initiation and independent from circRNA abundance. Thus, any analysis of the pathophysiological role of FTDP-17 mutants should not only rely on circRNA levels. Another unexpected finding was that the flanking introns dictate expression changes of circular RNAs in response to RNA editing. The prevailing view is that A>I editing weakens secondary structures [49] that promote backsplicing, which is expected to reduce circRNA formation. We observed the opposite effect for the 12➔10 circTau RNAs, indicating that A>I editing can have a positive effect on circRNA formation, depending on sequence context.

### 4.4. The 12➔7 circTau protein interacts with eIF4B

We unexpectedly found that the 12➔7 circTau protein binds to eIF4B through the DRYG domain of eIF4B. We observed this interaction in boiled cell lysates and did not detect other abundant proteins in the 12➔7 circTau immunoprecipitates, which suggests that this interaction is direct. It is specific for the 12->7 circTau protein as it was not seen for linear 0N4R tau and was much weaker for the 12➔10 circ proteins. EIF4B binding is a unique property of the 12➔7 circTau protein. It is worth noting that changes in translation are a hallmark of Alzheimer’s disease [50] and it remains to be determined whether 12➔7 circTau proteins contribute to this deregulation.

### 4.5 FTDP-17 mutation influence formation and translation of circular Tau RNAs

Our data suggest a complex interplay of FTDP-17 mutations with the formation and translation of circTau RNAs (**Figure 7**). The MAPT-pre mRNA adopts a secondary structure that promotes circRNA formation, likely by bringing backsplice sites into close proximity. The formation of a secondary pre-mRNA structure is promoted by repeat elements, which are often Alu-elements in primates [15]. CircRNAs form extended intermolecular base pairs and appear as rod-like structures in electron microscopy [17]. The base pairing generates double-stranded RNA regions that are substrates for ADAR-enzymes and multiple adenines of circTau RNAs are edited to inosines in the presence of ADAR activity [12]. The editing efficiency for a given adenine residue is less than 10%, i.e. lower than for known edited mRNAs [12]. It is possible that FTDP-17 mutations change the circRNA structure, which could promote backsplicing, and could thus explain our observed increase of circTau RNA formation for five FTDP-17 mutants. ADAR-activity is known to interfere with pre-mRNA secondary structures. In humans, the major target of ADAR are Alu-elements [51], which could interfere with the pairing of the backsplice sites. We found unexpectedly that this effect is sequence-context dependent, as ADAR1 reduced 12➔7 circTau RNA formation but increased 12➔10 circTau RNA formation. circTau RNAs are translated after being edited and the inosines most likely aid in ribosomal entry. All FTDP-17 mutants tested are translated into protein in the presence of ADAR1 and only three mutants, P301S, S350N and V363I slightly increase this effect. When ADAR variants are compared, ADAR1-p150 had by far the strongest effect on circRNA translation, independent of the FTDP-17 mutant present. This finding could have pathophysiological significance, as ADAR1-p150 is induced by cytokines that are elevated in AD [47,48]. We also identified three FTDP-17 mutants that are strongly translated by ADAR2, which could contribute to their protein expression.

**Figure 7:**
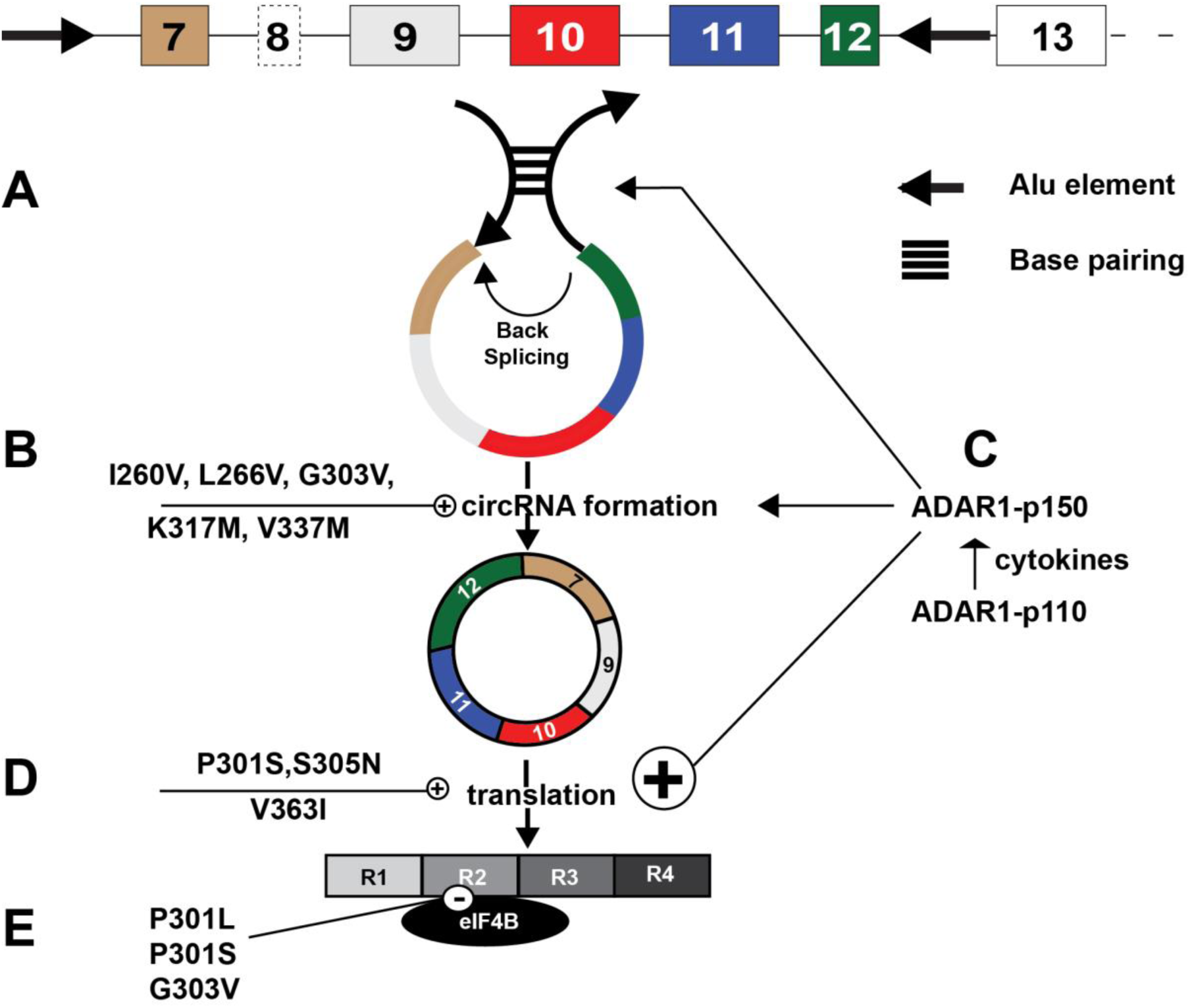
Model: FTDP-17 mutations influence circTau RNA processing and translation. **A.** Aided by Alu-elements (arrows), the MAPT pre-mRNAs adopt secondary structures that promote circRNA formation likely by bringing backsplice sites into close proximity. **B.** Several FTDP-17 mutations promote circRNA formation, possibly by changing the secondary structure. **C.** RNA editing influences circRNA processing depending on the sequence context. In the 12➔7 background, RNA editing reduces circRNA formation, for the 12➔10 background RNA editing increases circRNA formation. FTDP-17 mutations modulate the effect of RNA editing on circRNA formation. **D.** RNA editing strongly promotes circTau RNA translation and some FTDP-17 mutations further enhance circRNA translation. **E.** The 12➔7 circTau protein binds to eukaryotic initiation factor 4B, eIF4B, an interaction that could not be detected with linear tau protein.

Finally, we found that the 12➔7 circTau protein binds to eIF4B, which is a feature distinct from linear tau protein. FTDP-17 mutations located in the second microtubule repeat binding domain reduce this interaction and could thus interfere with its function in translational initiation that needs to be determined.

## Conclusion

FTDP-17 mutations influence circTau RNA and protein expression, as well as the interaction between circTau protein and eukaryotic initiation factor 4B. Circular Tau RNAs should be considered, when determining the pathophysiological effect of FTDP-17 mutations.

## Acknowledgments

This work was supported by the Department of Defense (W81XWH-19-1-0502), the National Institutes of Health (1R21AG067438-0) and National Science Foundation (# 2221921). We thank the Cedars-Sinai proteomics and metabolomics core for access to mass spectrometers.

## Supplemental Figures

**Supplemental Figure 1.**
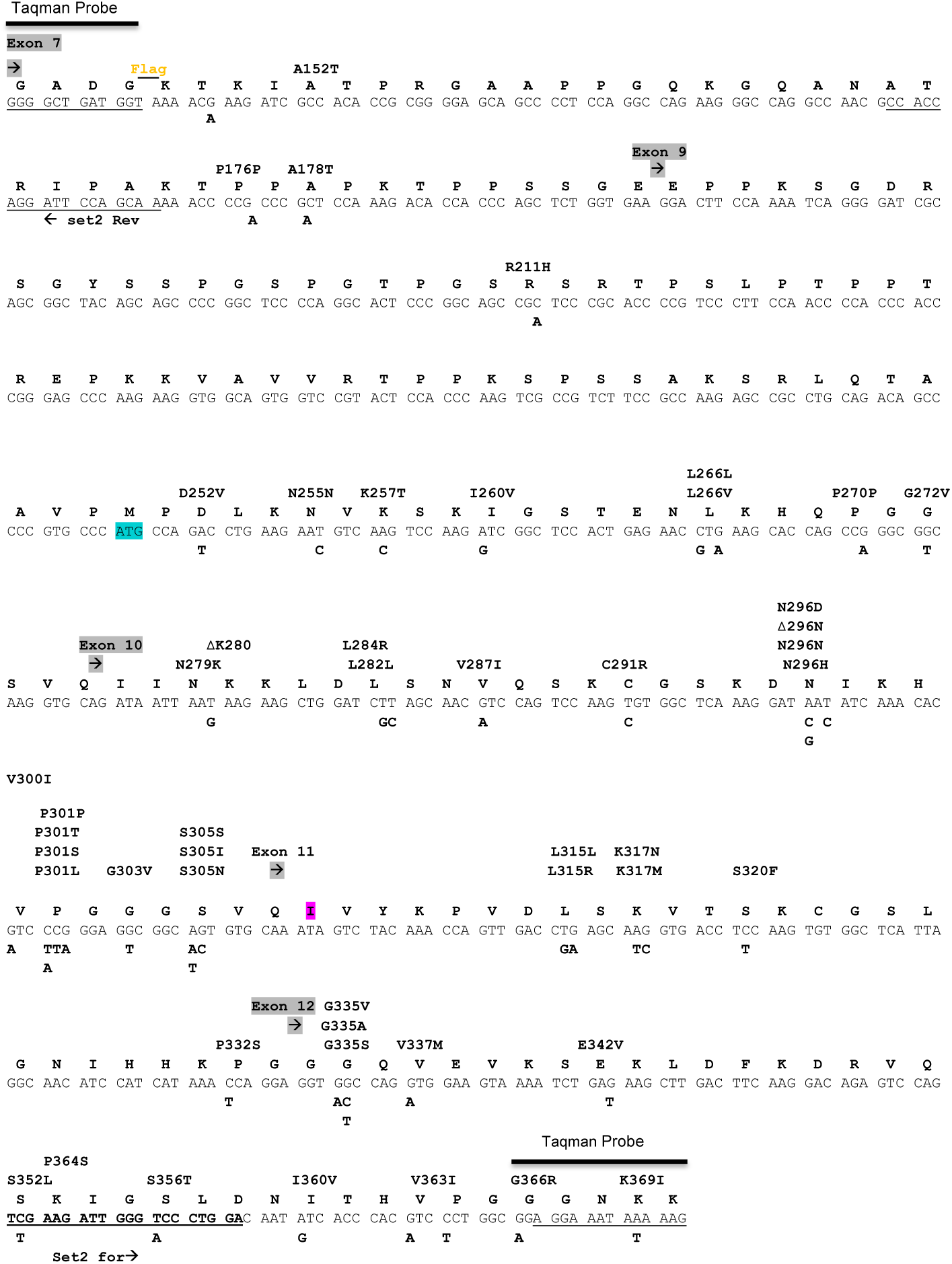
The sequence of the 12➔7 circTau RNA is shown. The location of the TaqMan probe, and the forward and reverse primers are indicated. FTDP-17 mutations are indicated on top of the sequence and the corresponding nucleotide changes are shown below.

**Supplemental Figure 2.**
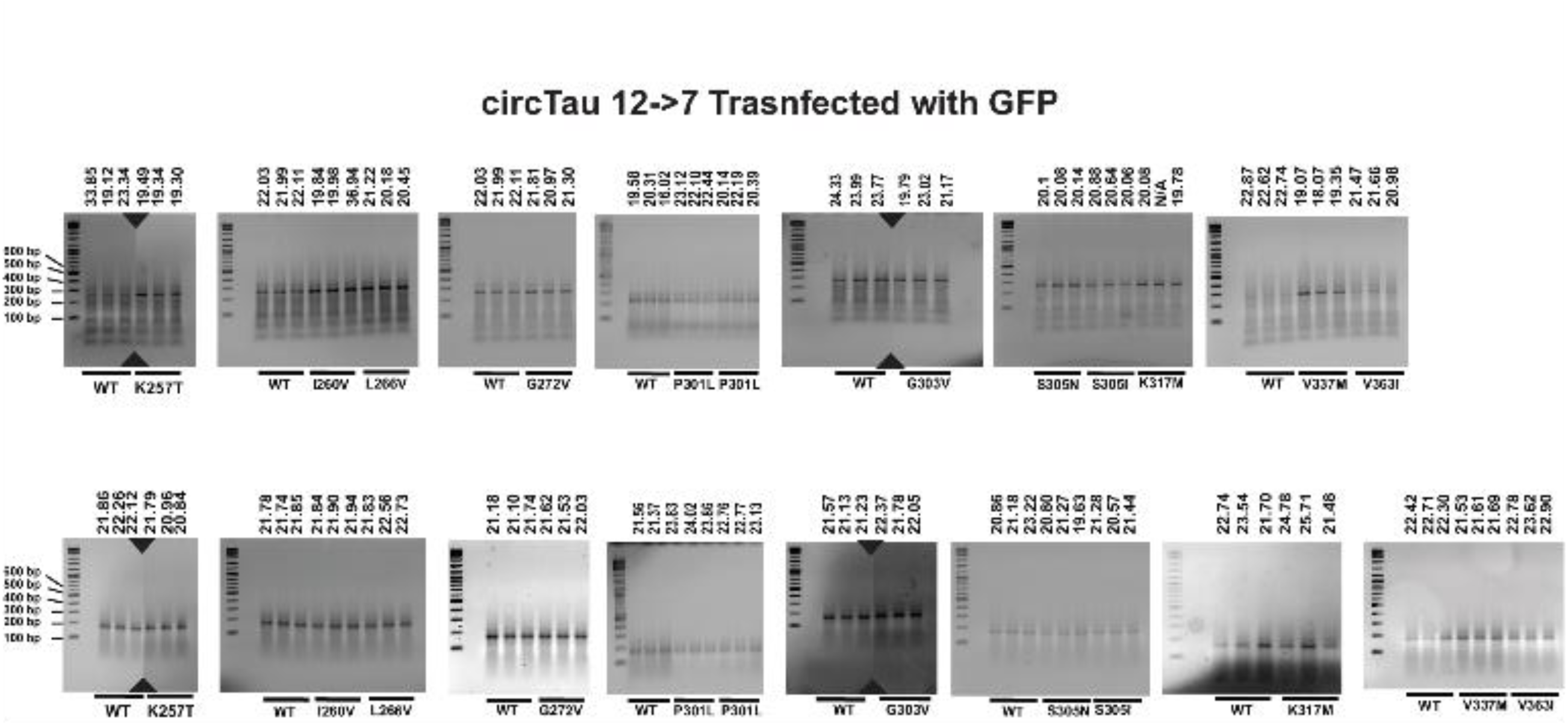
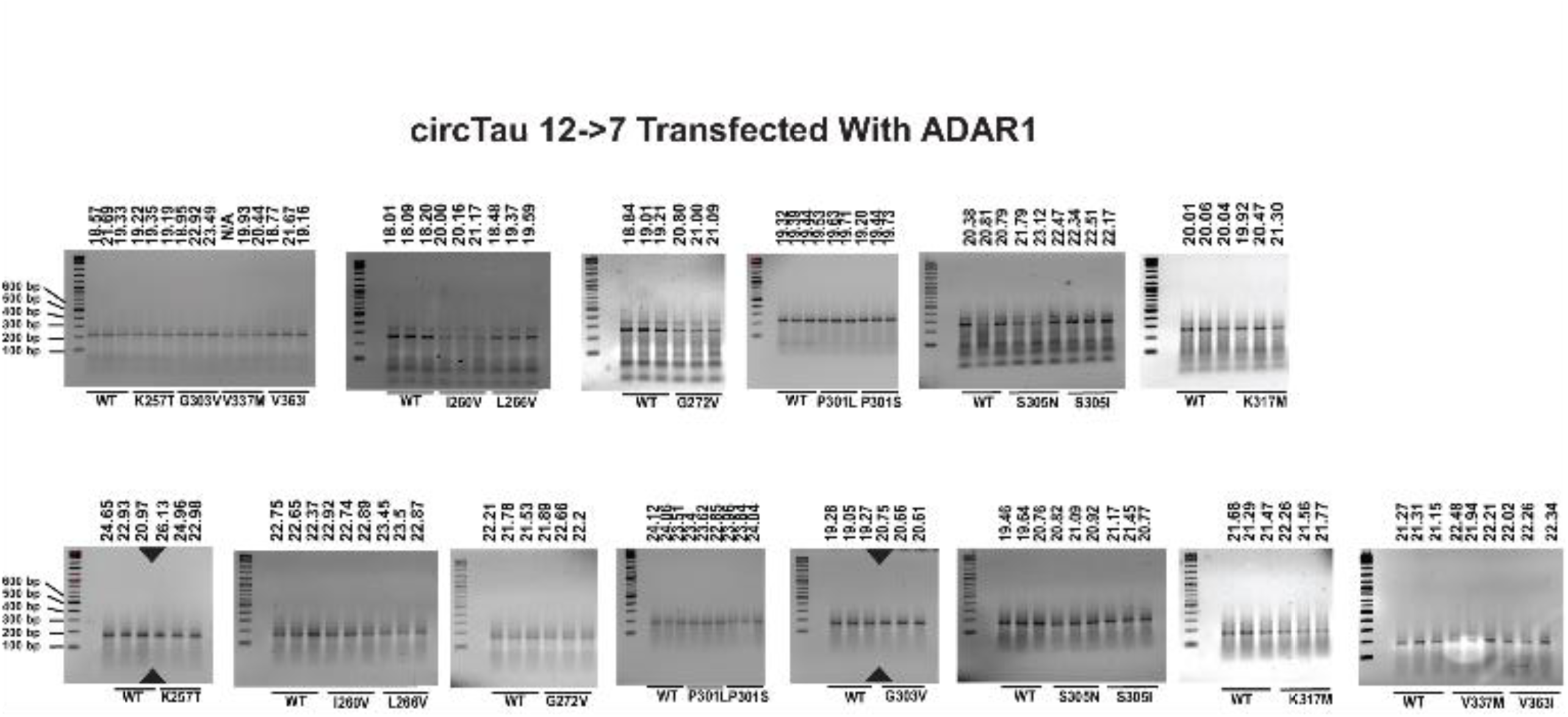
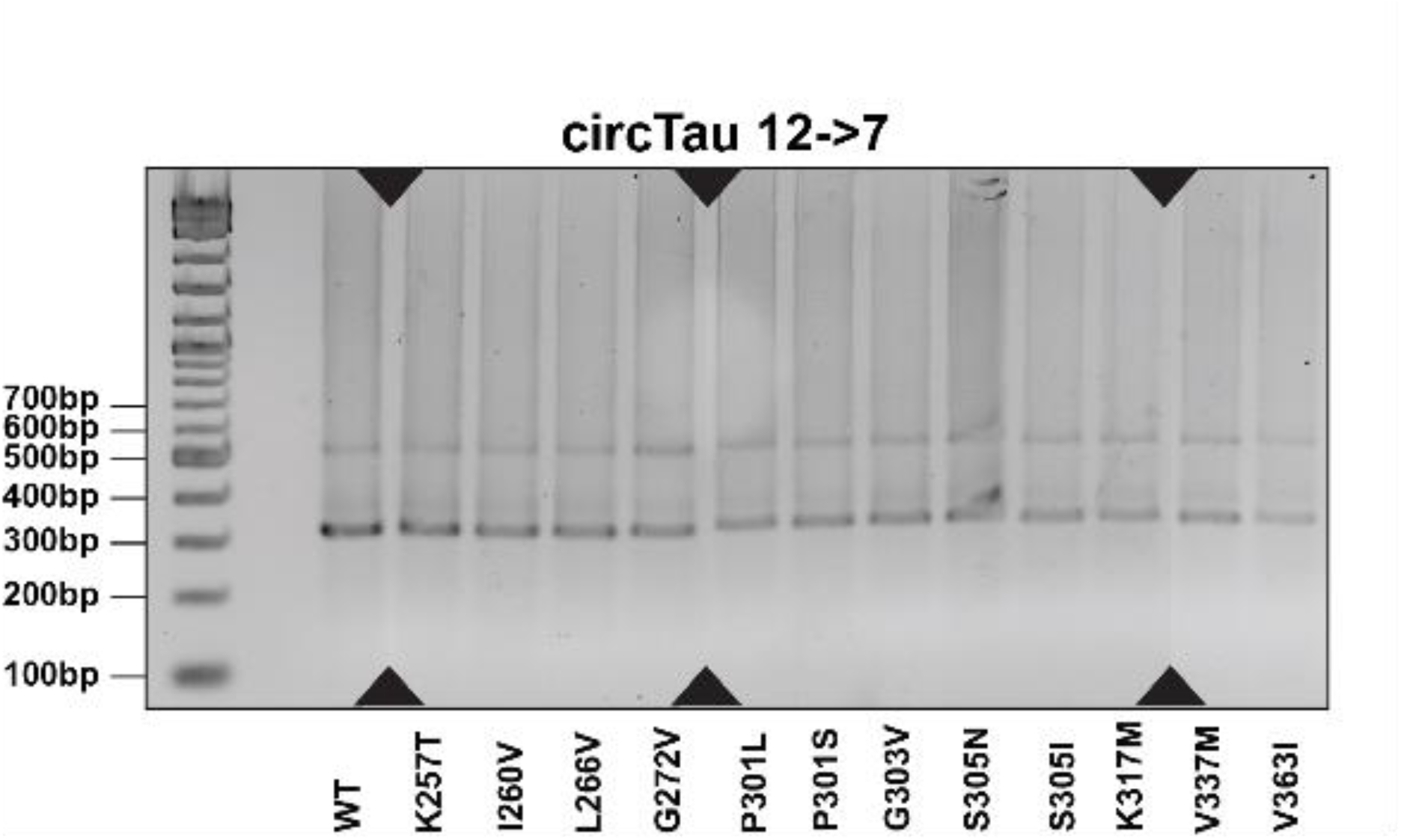
**A.** Gel electrophoresis of the end points of the TaqMan RT-PCR reactions. 12➔7 expression constructs co-transfected with GFP were used. The ct values are shown on top of the gels. The expected band is 216 nt. The mutants are indicated. The top row shows detection of the circTau RNA, the bottom row shows the detections of GAPDH. **B.** The experiments were similar to (A), but RNAs from HEK293T cells cotransfected with ADAR1-p150 were used. **C.** Agarose gel of the RT-PCR results for circTau RNA after 40 cycles. The expected size is 315 nt and the second weaker band around 500 nt likely represents a rolling circle reverse transcription. Primers used were circdetect Fw2: AATAAGAAGCTGGATCTTAGC and circdetect rev2: GATCTTTATAATCACCGTCAT

**Supplemental Figure 3.**
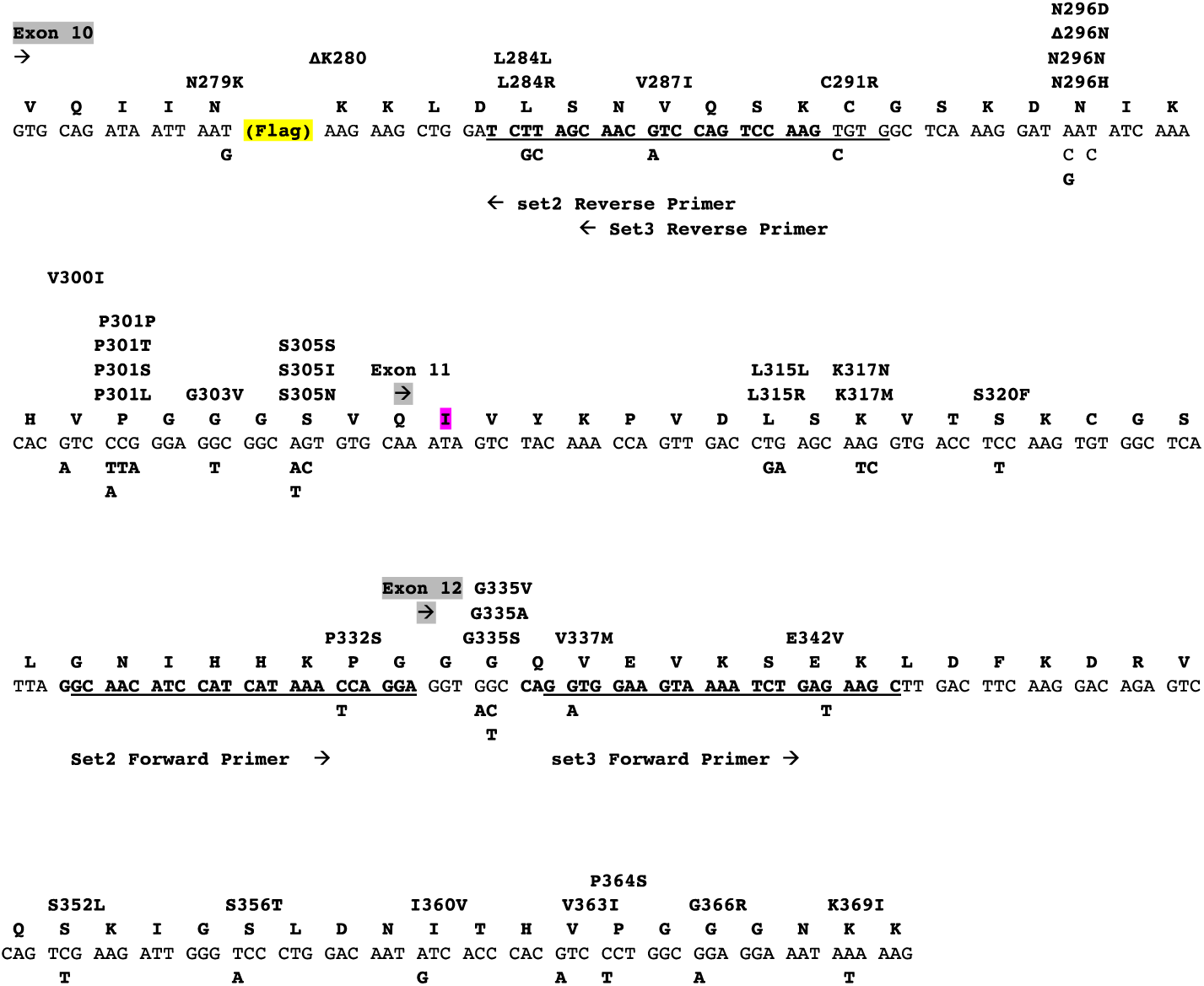
The sequence of the 12➔10 circTau RNA is shown. The location of the forward and reverse primers is indicated. FTDP-17 mutations are indicated on top of the sequence and the corresponding nucleotide changes are shown below.

**Supplemental Figure 4.**
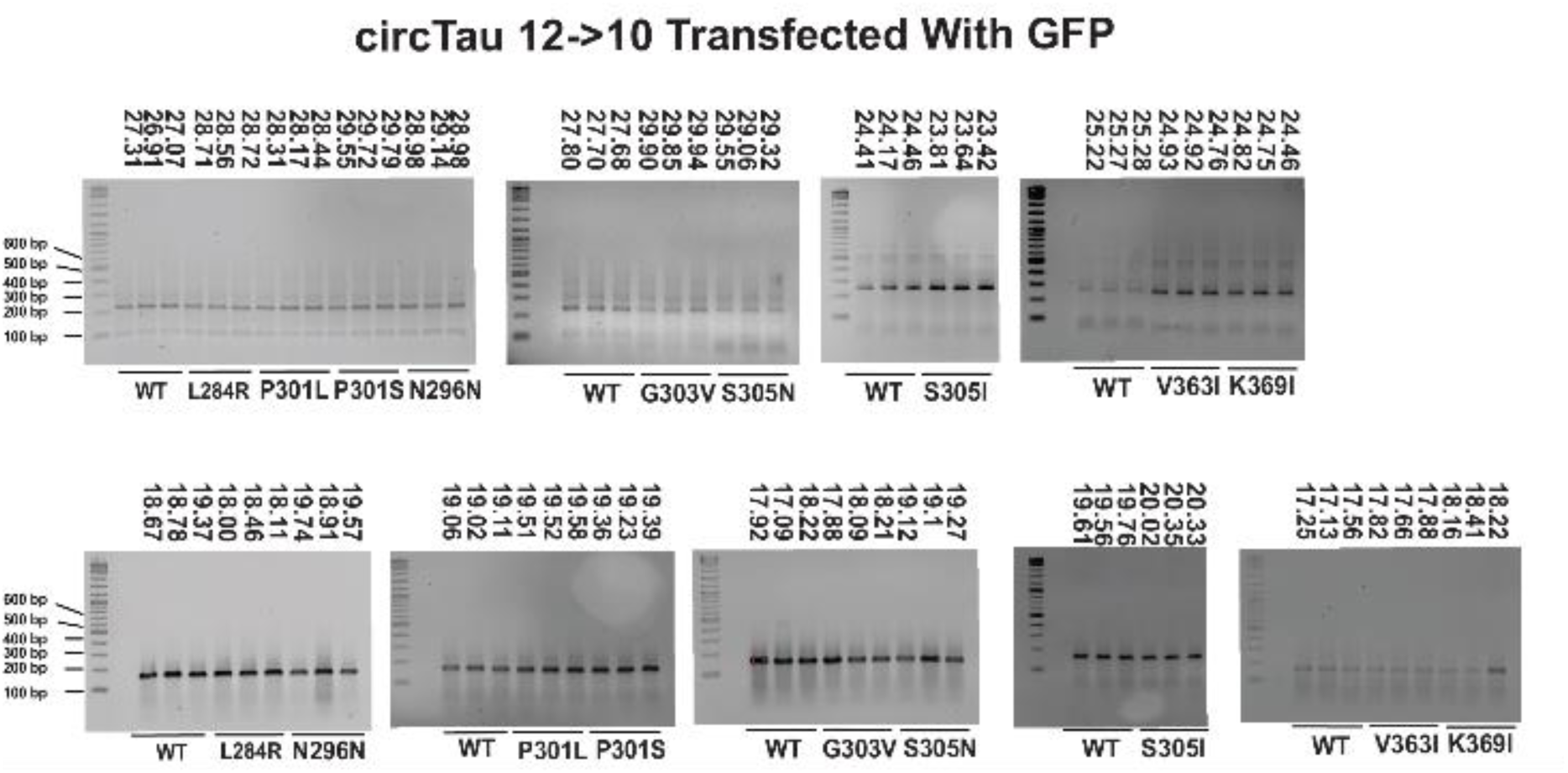
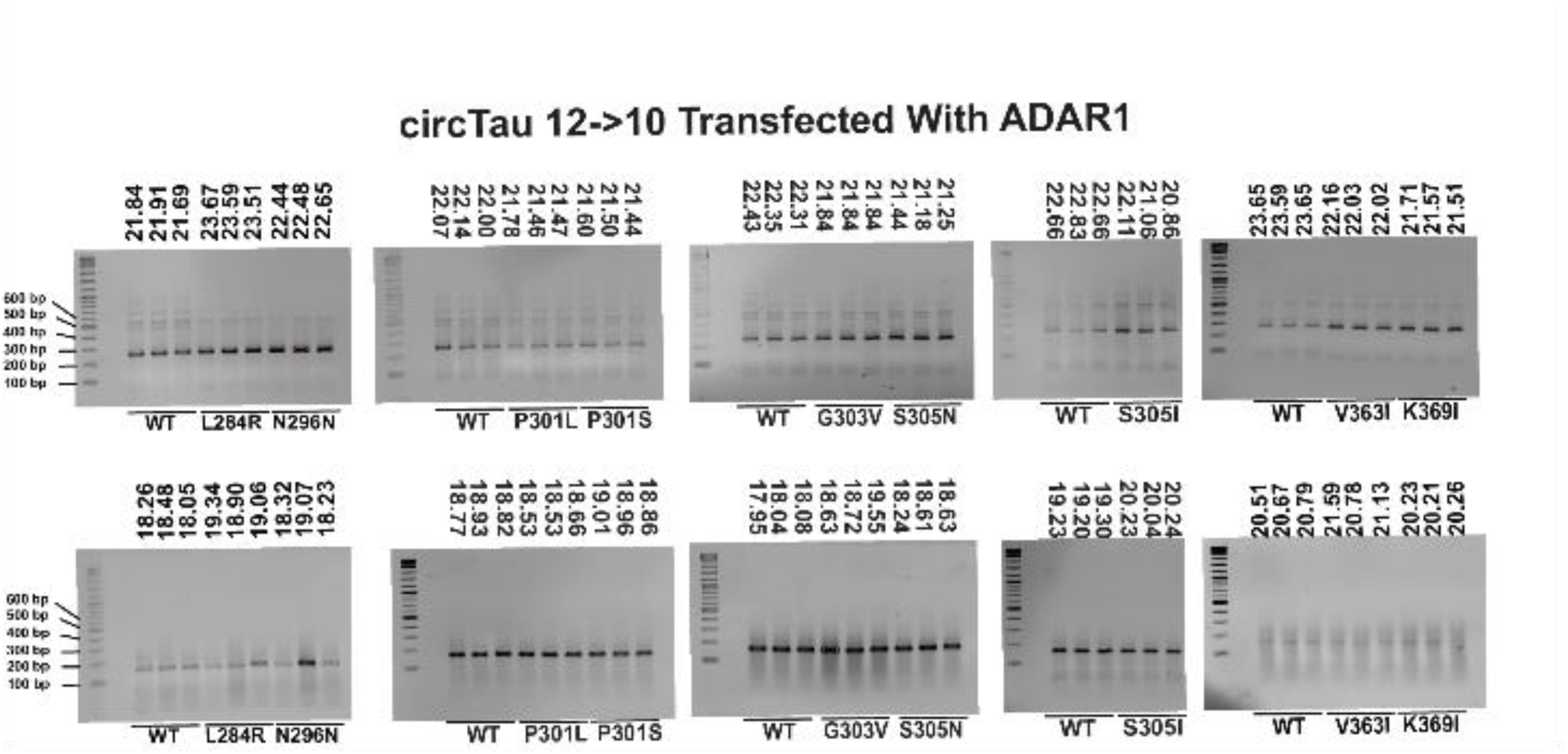
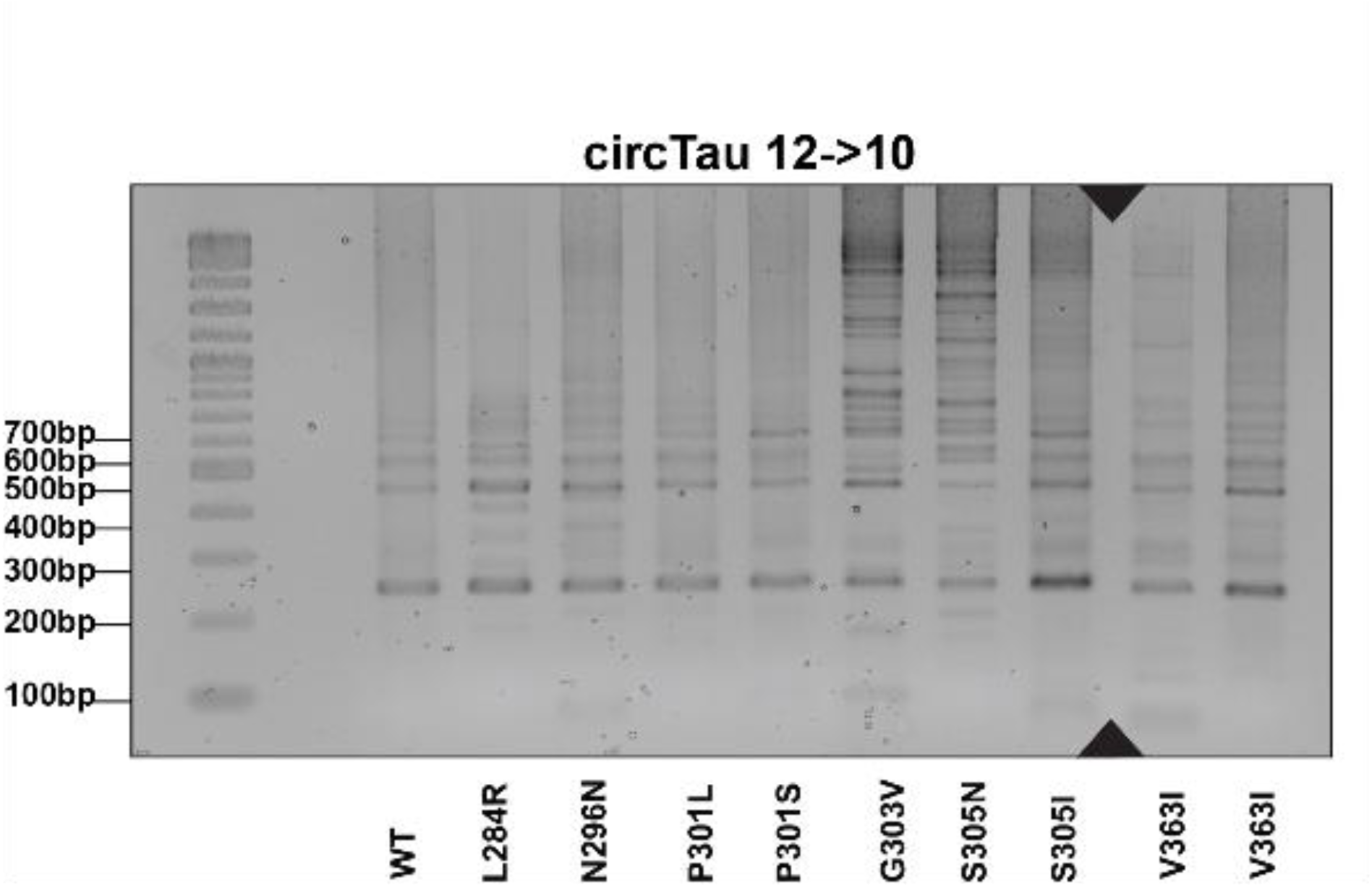
**A.** Gel electrophoresis of the end points of the SYBR green reaction RT-PCR reactions. 12➔10 expression constructs co-transfected with GFP were used. The ct values are shown on top of the gels. The expected band is 246 nt. The mutants are indicated. The top row shows detection of the circTau RNA, the bottom row shows the detections of GAPDH using a TaqMan assay. **B.** The experiments were similar to (A), but RNAs from HEK293T cells cotransfected with ADAR1-p150 were used. **C.** Agarose gel of the RT-PCR results for circTau RNA after 40 cycles. The expected size is 246 nt and the second weaker band of about 400 nt likely represents a rolling circle reverse transcription.

**Supplemental Figure 5.**
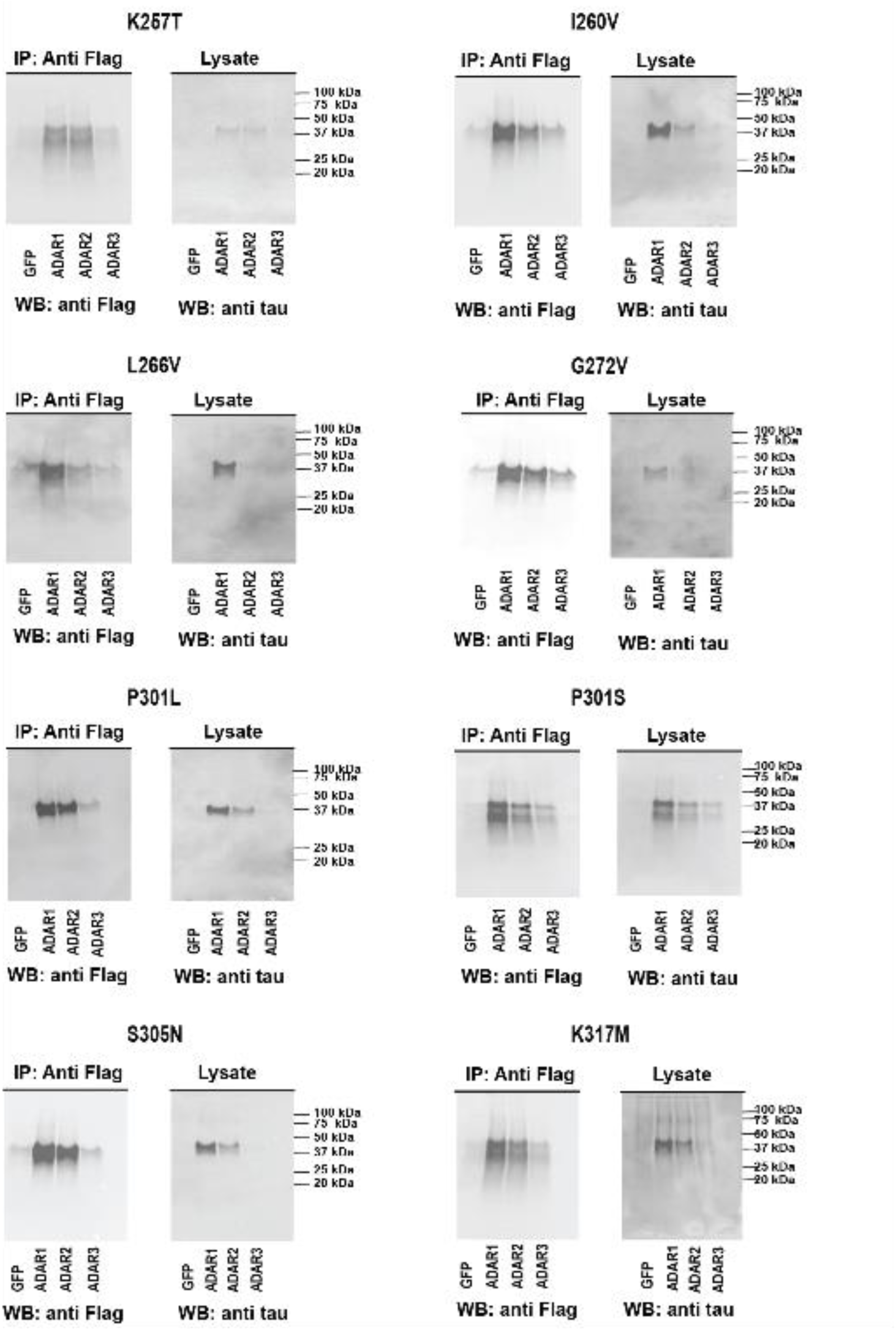
Expression clones of the FTDP-17 mutants in the 12➔7 circTau RNA background were cotransfected with equal amounts of expression clones for EGFP, ADAR1-p150, ADAR2 and ADAR3 and Immunoprecipitated using anti-Flag. The immunoprecipitates were analyzed by Western blot using anti Flag and anti tau antisera. The analysis of the lysates with anti Flag is shown in Figure 5.

**Supplemental Figure 6.**
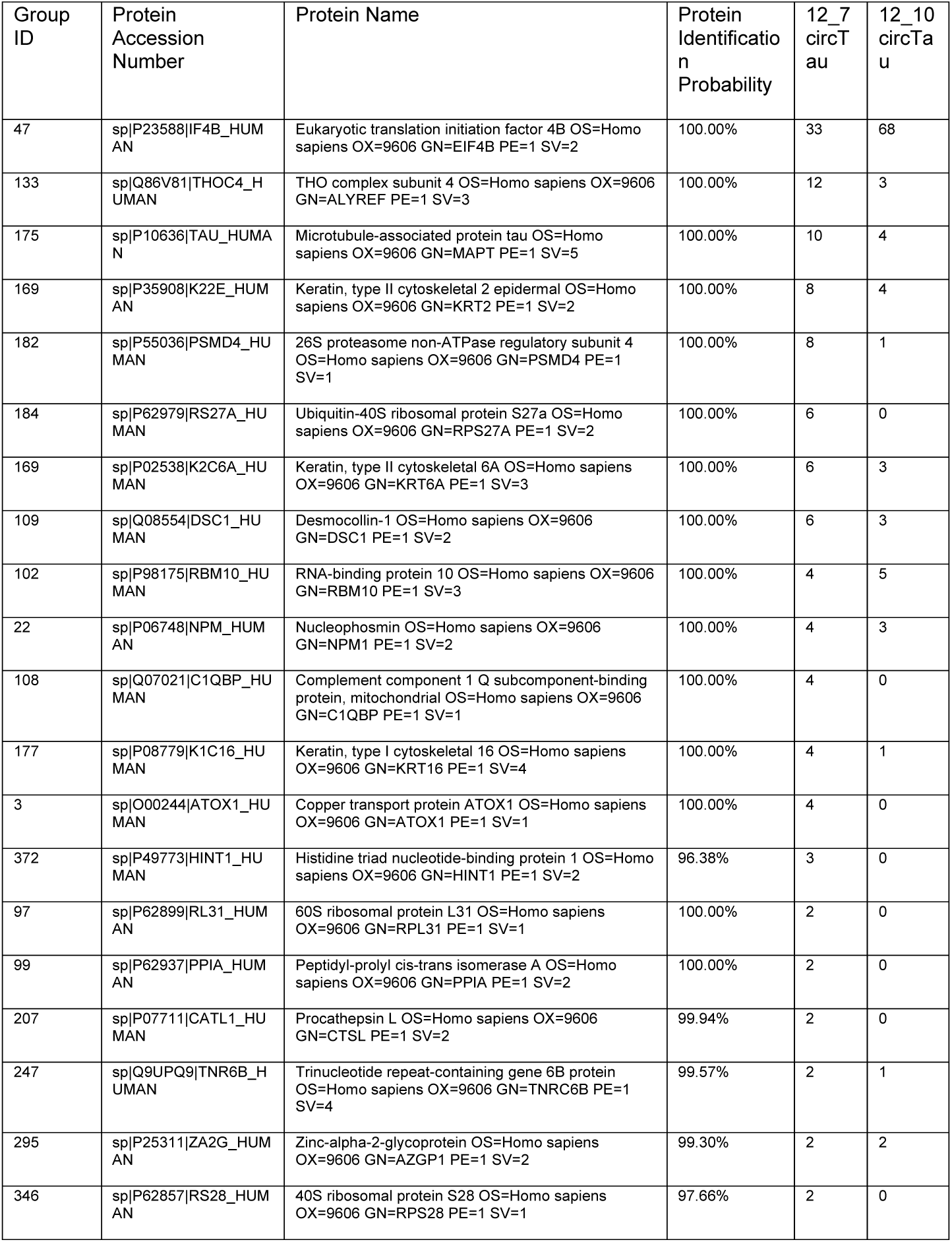
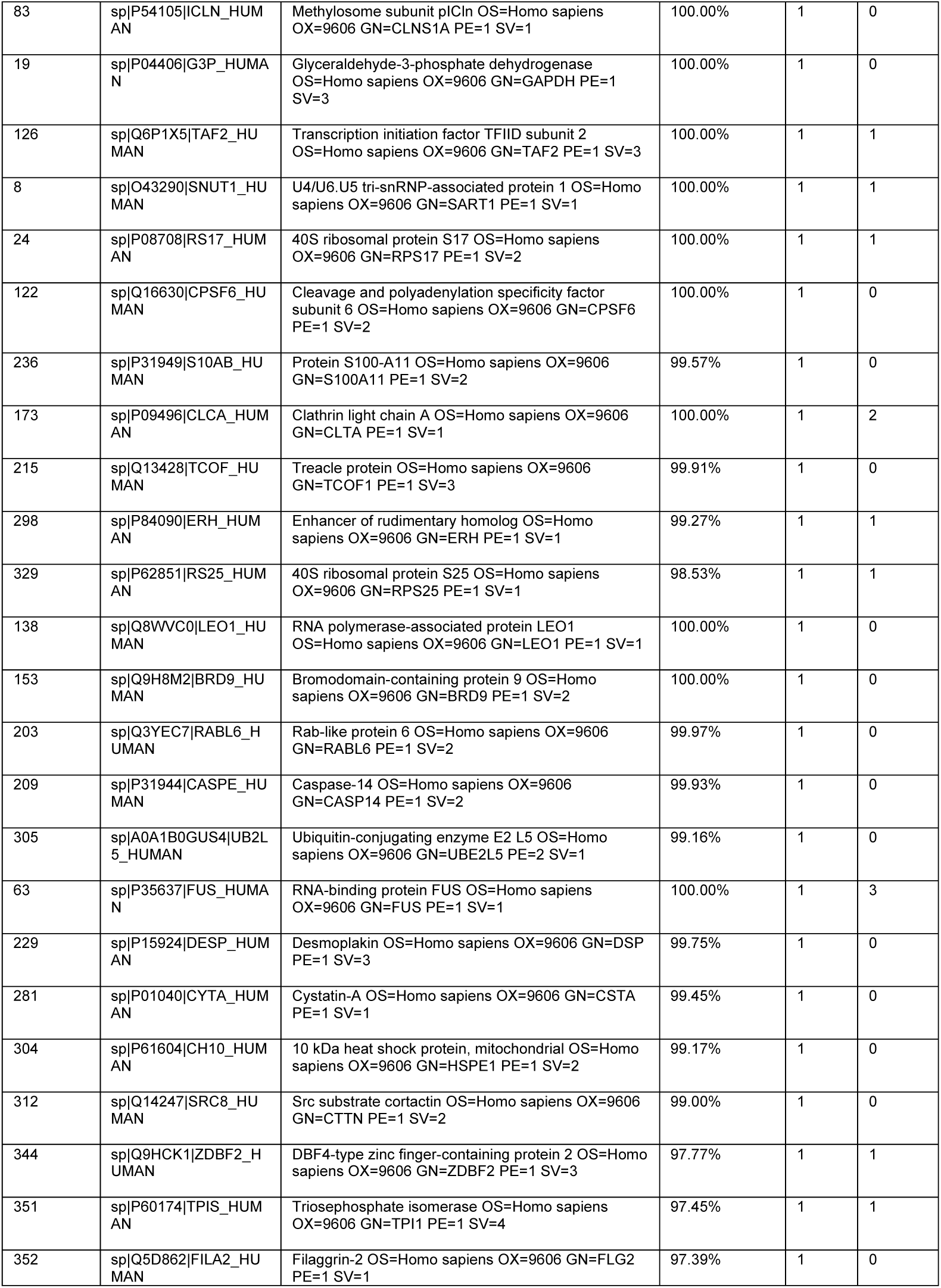
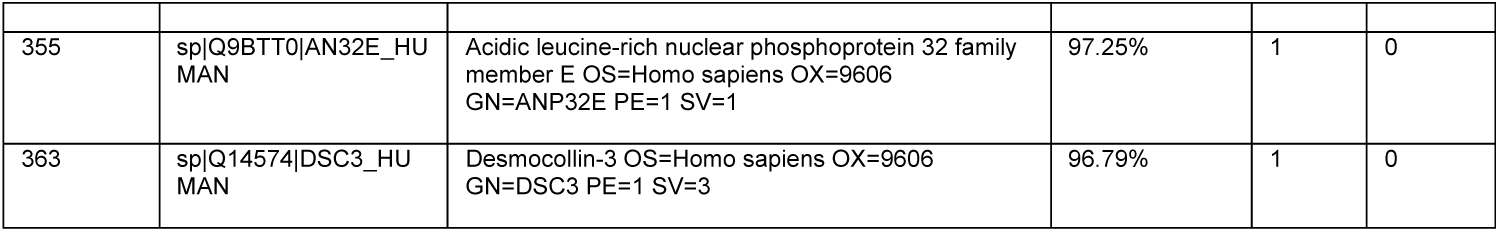
Proteins interacting with circTau. 12➔10 and 12➔7 circTau expression clones were transfected into HEK293T cells and immunoprecipitates analyzed using mass-spectroscopy after the total immunoprecipitate was digested.

**Supplemental Figure 7.**
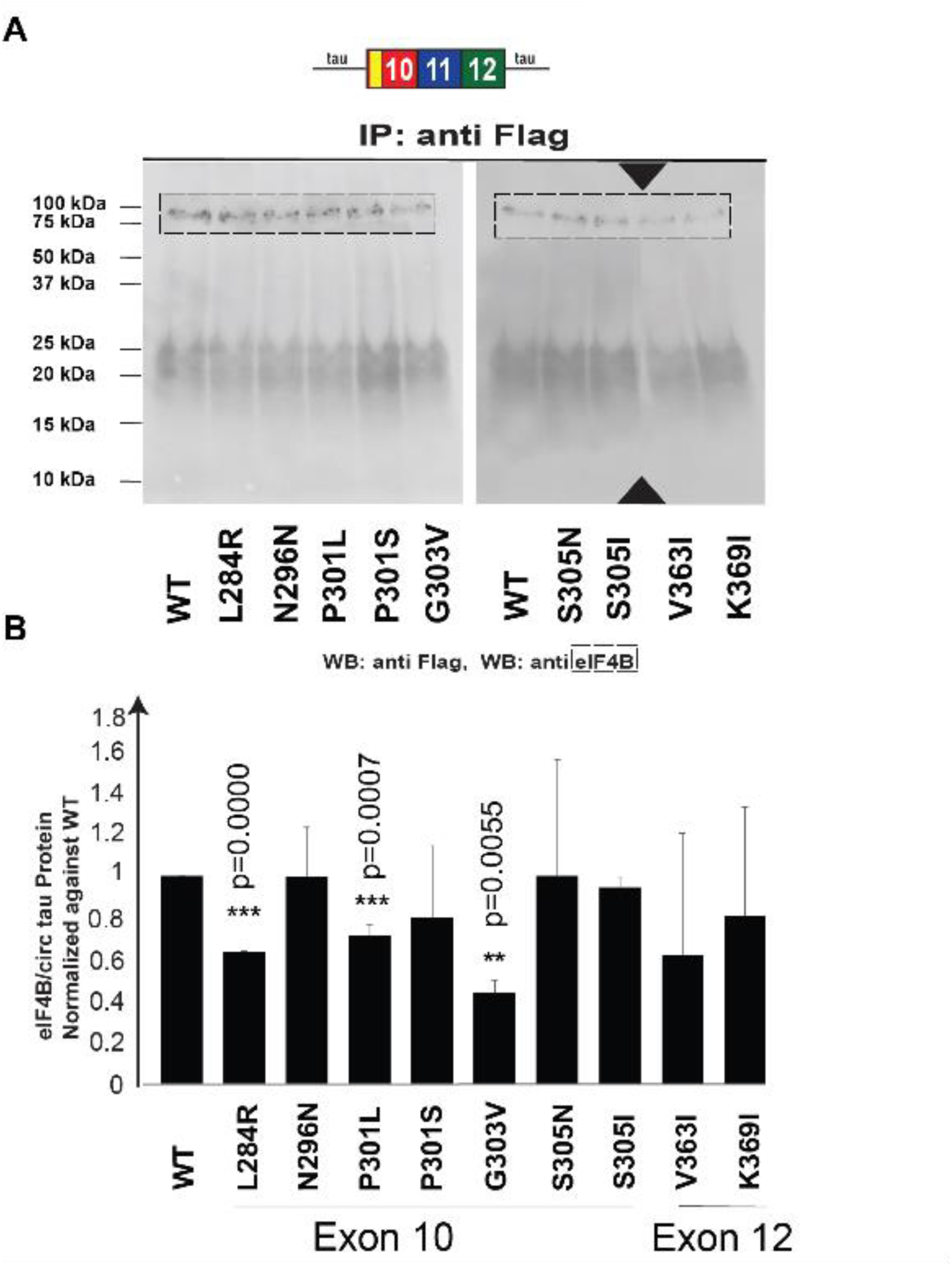
**A.** The 12➔10 circTau mutants indicated were expressed in HEK293T cells and immunoprecipitated with anti-Flag. Protein was detected in Western Blot using anti-Flag. The immunoprecipitates were probed with eIF4B antibody for endogenous eIF4B binding. **B.** Quantification of at least three independent experiments.

